# Covalent inhibition of endoplasmic reticulum chaperone GRP78 disconnects the transduction of ER stress signals to inflammation and lipid accumulation in diet-induced obese mice

**DOI:** 10.1101/2021.08.20.457085

**Authors:** Dan Luo, Ni Fan, Xiuying Zhang, Fung Yin Ngo, Jia Zhao, Wei Zhao, Min Huang, Ding Li, Yu Wang, Jianhui Rong

## Abstract

Targeting endoplasmic reticulum (ER) stress, inflammation and metabolic dysfunctions may halt the pathogenesis of obesity and thereby reduce the prevalence of diabetes, cardiovascular disesases and cancers. The present study was designed to elucidate the mechnaisms by which plant-derived celastrol ameliorated inflammation and lipid accumulation in obesity. The mouse model of diet-induced obesity was induced by feeding high-fat diet for 3 months and subsequently intervented with celastrol for 21 days. Hepatic and adipose tissues were analysed for lipid accumulation, macrophage activation and biomarker expression. As result, celastrol effectively reduced body weight, suppressed ER stress, inflammation and lipogenesis while promoted hepatic lipolysis. RNA-sequencing revealed that celastrol-loaded nanomicelles restored the expression of 49 genes that regulate ER stress, inflammation and lipid metabolism. On the other hand, celastrol-PEG4-alkyne was synthesized for identifying celastrol-bound proteins in RAW264.7 macrophages. ER chaperone GRP78 was identified by proteomics approach for celastrol binding to the residue Cys^41^. Upon binding and conjugation, celastrol diminished the chaperone activity of GRP78 by 130-fold and reduced ER stress in palmitate-challenged cells, while celastrol analogue lacking quinone methide failed to exhibit anti-obesity effects. Thus, covalent GRP78 inhibition may induce the reprograming of ER signaling, inflammation and metabolism against diet-induced obesity.

## Introduction

Dysregulation of cross-talks between endoplasmic reticulum (ER) stress, inflammation and metabolic pathways hallmark the pathogenesis of obesity, diabetes, atherosclerosis, hypertension and acute myocardial infarction (Van Gaal et al. 2006, Rocha and Libby 2009, Flegal et al. 2010). The 78-kDa glucose-regulated protein (GRP78) is an integral ER stress sensor and holds inositol-requiring enzyme 1 (IRE1), protein kinase R-like ER kinase (PERK) and activating transcription factor 6 (ATF6) in the ER lumen to prevent the activation of corresponding down-stream substrates under the physiological conditions (Szegezdi et al. 2006, Cnop et al. 2012). Upon the challenge by stress stimuli, however, unfolded or misfolded proteins are overwhelmingly accumulated in ER compartment, disrupt the complexes of GRP78 with IRE1, PERK and ATF6, and ultimately induce unfolded protein response (UPR) (Cnop et al. 2012). Obesity and high-fat diet (HFD) not only induce persistent ER stress, leptin resistance and insulin resistance via activating protein kinase JNK and inducing the phosphorylation of insulin receptor substrate 1 (IRS-1) (Ozcan et al. 2004, Ozcan et al. 2009), but also promote the generation of unfolded or misfolded proteins and the activation of proinflammatory signaling pathways (Hotamisligil 2010, Pagliassotti et al. 2016). On the other hand, hyperactive ER stress drives the pro-inflammatory M1 polarization of adipose tissue macrophages (ATMs) and exacerbates inflammation and metabolic dysfunctions in obesity (Bujisic and Martinon 2017, Shan et al. 2017). Importantly, GRP78 is essential for the differentiation of preadipocytes into adipocytes, indicating a critical role in the control of fat contents in body (Zhu et al. 2013). The activation of ER stress pathways (e.g., PERK and eIF2α) inversely induces the upregulation of GRP78 expression (Ozcan et al. 2004, Shan et al. 2017). Thus, GRP78 may be an important molecular target for the development of new therapies against obesity, type 2 diabetes, atherosclerosis, hypertension and acute myocardial infarction.

Plant-derived pentacyclic triterpene celastrol is recently identified as the best anti-obesity drug candidate (Liu et al. 2015). It is now known that celastrol may induce rapid loss of body weight through several mechanisms as follows: 1) To enhance leptin activity and reduce food intake in obese mice (Greenhill 2015, Liu et al. 2015); 2) To improve insulin sensitivity via inhibiting NF-kB pathway and control the progression of obesity via enhancing antioxidant capacity and lipid metabolism (Kim et al. 2013, Wang et al. 2014); 3) To activate HSF1-PGC1α transcriptional axis and elicit beneficial metabolic changes against obesity (Ma et al. 2015). At the cell level, celastrol could effectively suppress proinflammatory M1 macrophage polarization and enhance anti-inflammatory M2 macrophage polarization (Luo et al. 2017). To improve the bioavailability, hydrophobic celastrol was loaded into PEG-PCL nano-micelles to yield Nano-celastrol with highly similar anti-obesity effects and largely reduced gastrointestinal issues (Zhao et al. 2019). However, the primary molecular target still remains elusive, thereby limiting the development of novel anti-obesity drugs.

The aim of the present study is to discover the protein targets for celastrol and determine the impact of covalent celastrol-protein conjugation on ER stress in diet-induced obesity. We synthesized celastrol-PEG4-alkyne bearing an alkyne (-C≡C-) group as a molecular probe for the affinity isolation of celastrol-bound proteins. We further determined the *in vitro* and *in vivo* effects of celastrol on ER stress, inflammation and lipid metabolism.

## Results

### Celastrol diminished lipid accumulation in diet-induced obesity

To examine the effects of celastrol on lipid metabolism, we visualized the morphology of adipocytes and the profiles of fatty acids in livers and adipose tissues from diet-induced obese mice. Firstly, liver tissues and adipose tissues were examined by H&E staining. As shown in Figure 1A and 1B, celastrol effectively ameliorated adipose hypertrophy and lipid accumulation in both livers and epididymal fat pads. Secondly, the compositions of fatty acids in liver and adipose tissues were profiled by GC-MS technology. Figure 1C and 1D showed the GC-MS chromatograms of lipid extracts from the livers and adipose tissues of three treatment groups. Based on the quantitative analysis in Figure 1E and 1F, HFD elevated the contents of several polyunsaturated fatty acids (e.g., C16:1, trans-C18:1, cis-C18:1, C18:2, C18:3, cis-C20:2, cis-C20:3) whereas celastrol ameliorated HFD-induced elevations of specific fatty acids in liver (Figure 1C and 1E). By contrast, HFD decreased the contents of long chain polyunsaturated fatty acids eicosoids (e.g., cis-C20:1, cis-C20:3, cis-C22:6) in epididymal fat pads while celastrol showed little activity (Figure 1D and 1F).

**Figure 1.**
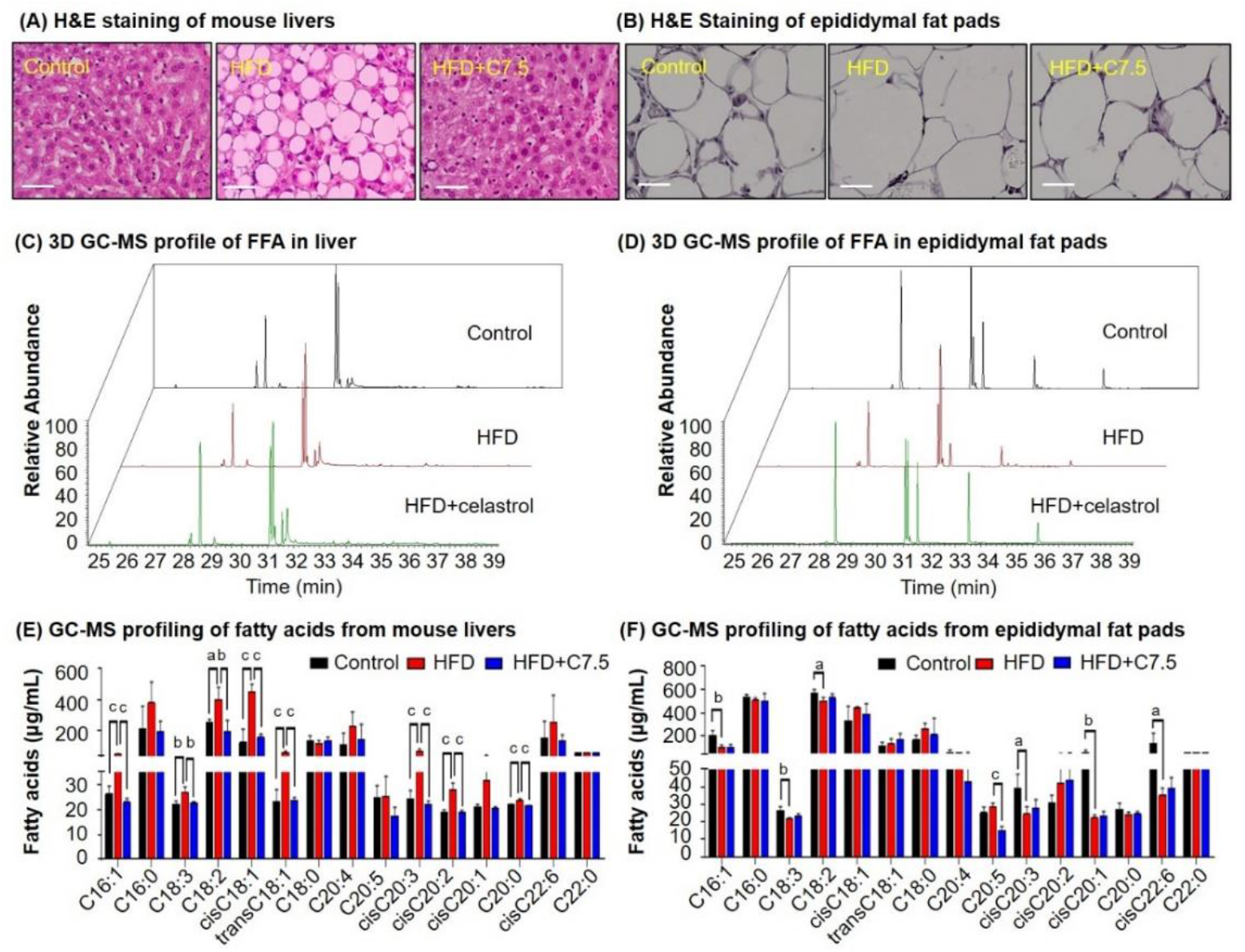
Celastrol ameliorates adipose hypertrophy and lipid metabolism. (A & B) H&E staining of mouse hepatic tissues and epididymal fat pads. After 21-day treatment with vehicle or celastrol, the livers and epididymal fat pads were recovered from four groups of C57/BL6 mice (i.e., Control, HFD, HFD+C5, HFD+C7.5) and stained with H&E stain. Representative images were shown. Scale bar represented 34 μm in length. (C & D) 3-D GC-MS chromatograms of free fatty acids from murine livers and epididymal fat pads. After 21-day treatment with vehicle or celastrol, livers and epididymal fat pads were recovered from four groups of C57/BL6 mice (i.e., Control, HFD, HFD+C5, HFD+C7.5) for fatty acids extraction and GC-MS profiling. (E & F) GC-MS Profiling of fatty acids from murine livers and epididymal fat pads. Free and conjugated fatty acids were profiled by GC-MS. Total amounts of individual fatty acids were quantified. The results were presented as mean ± SD of three independent experiments. a, *p*<0.05; b, *p*<0.01; c, *p*<0.001. **Source data 1.** Data for Figure 1A-1D. **Source data 2.** Data for Figure 1E-1F.

### Celastrol inhibited lipogenesis while promoted lipolysis and thermogenesis

To investigate how celastrol regulates lipid metabolism, we employed qRT-PCR technique to determine the expression of the representative biomarkers in lipogenesis, lipolysis and thermogenesis. As shown in Figure 2A, HFD upregulated the expression of adipogenic genes (e.g., PPAR-γ, C/EBP-α, aP2) and adipocyte-secreted hormones (e.g., resistin, adipsin) while suppressed the lipolytic genes (e.g., CPT-1, CAT, ACO) and thermogenic gene UCP2 in liver. After the treatment at the dose of 7.5 mg·kg^-1^·d^-1^ for 7-days, celastrol effectively down-regulated the expression of the adipocyte-secreted hormones (e.g., resistin, adipsin), restored the lipolytic genes (e.g., CPT-1, CAT) and thermogenic gene UCP2 and decreased the expression of the selected adipogenic genes (e.g., PPAR-γ, C/EBP-α, aP2) to the levels below control. After the animals were treated with celastrol at the doses of 5.0 and 7.5 mg·kg^-1^·d^-1^ for 21 days, the levels of these biomarkers in liver and adipose tissues were determined by qRT-PCR technique. As shown in Figure 2B, celastrol effectively decreased the upregulation of the lipogenic genes (e.g., PPAR-γ, C/EBP-α, aP2) and adipocyte-secreted hormones (e.g., leptin, resistin, adipisin) in livers against HFD stimulation. By contrast, celastrol upregulated the lipolytic genes (e.g., CPT-1, CAT, ACO) and thermogenic gene UCP2 in liver in a dose-dependent manner whereas HFD did not show any effects. As shown in Figure 2C, in adipose tissues, celastrol at the dose of 7.5 mg·kg^-1^·d^-1^ attenuated the upregulation of the selected lipogenic genes and adipocyte-secreted hormones against HFD stimulation. On the other hand, both HFD and celastrol did not show much effects on the expression of the selected lipolytic and thermogenic genes.

**Figure 2.**
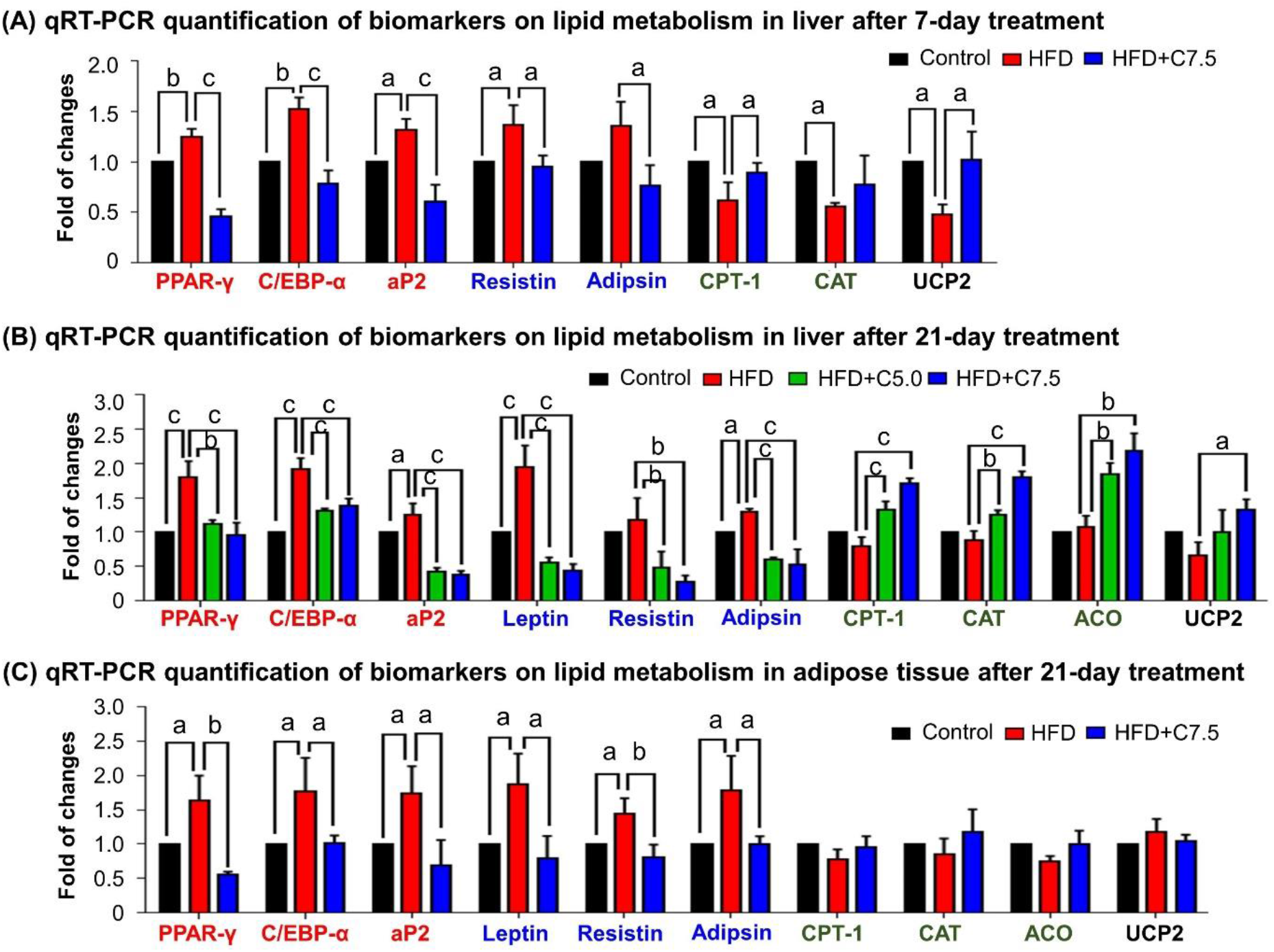
Celastrol restores the expression of the biomarkers in lipid metabolism in livers and epididymal adipose tissues. (A) One-week effects of celastrol on the biomarkers of lipid metabolism in liver tissues. After 7-day treatment, total RNAs were extracted from livers and analyzed by qRT-PCR technique using Qiagen QuantiTect SYBR Green PCR Kit and specific DNA primers. N = 3. (B) Three-week effects of celastrol on the biomarkers in lipid metabolism in liver tissues. After 21-day treatment, total RNAs were extracted from livers and analyzed by qRT-PCR technique using Qiagen QuantiTect SYBR Green PCR Kit and specific DNA primers. N = 3. (C) Three-week effects of celastrol the biomarkers in lipid metabolism in epididymal adipose tissues. After 21-day treatment, total RNAs were extracted from livers and analyzed by qRT-PCR technique using Qiagen QuantiTect SYBR Green PCR Kit and specific DNA primers. N = 3. HFD, HFD only; C5.0, celastrol (5 mg·kg^-1^·d^-1^); C7.5, celastrol (7.5 mg·kg^-1^·d^-1^); a, *p*<0.05; b, *p*<0.01; c, *p*<0.001. **Source data.** Data for Figure 2 A-C.

### Celastrol decreased ER stress biomarker GRP78 in hepatic and adipose tissue macrophages

To validate the *in vivo* effects of celastrol on ER stress, we employed immunofluorescence staining to examine the expression of biomarker GRP78 in hepatic and adipose tissue macrophages from diet-induced obese mice. Mouse liver and epididymal fat pads were recovered from three groups of animals (i.e., Control, HFD, HFD+Celastrol (7.5 mg·kg^-1^·d^-1^)), and stained with antibodies against GRP78, whereas macrophages were identified with antibody against biomarker CD68 and the cell nuclei were stained with DAPI. As shown in Figure 3A and 3B, macrophage infiltration and GRP78 expression were greatly enhanced in liver and adipose tissues in obese mice, whereas celastrol treatment effectively inhibited macrophage infiltration and reduced the protein level of GRP78 in macrophages.

**Figure 3.**
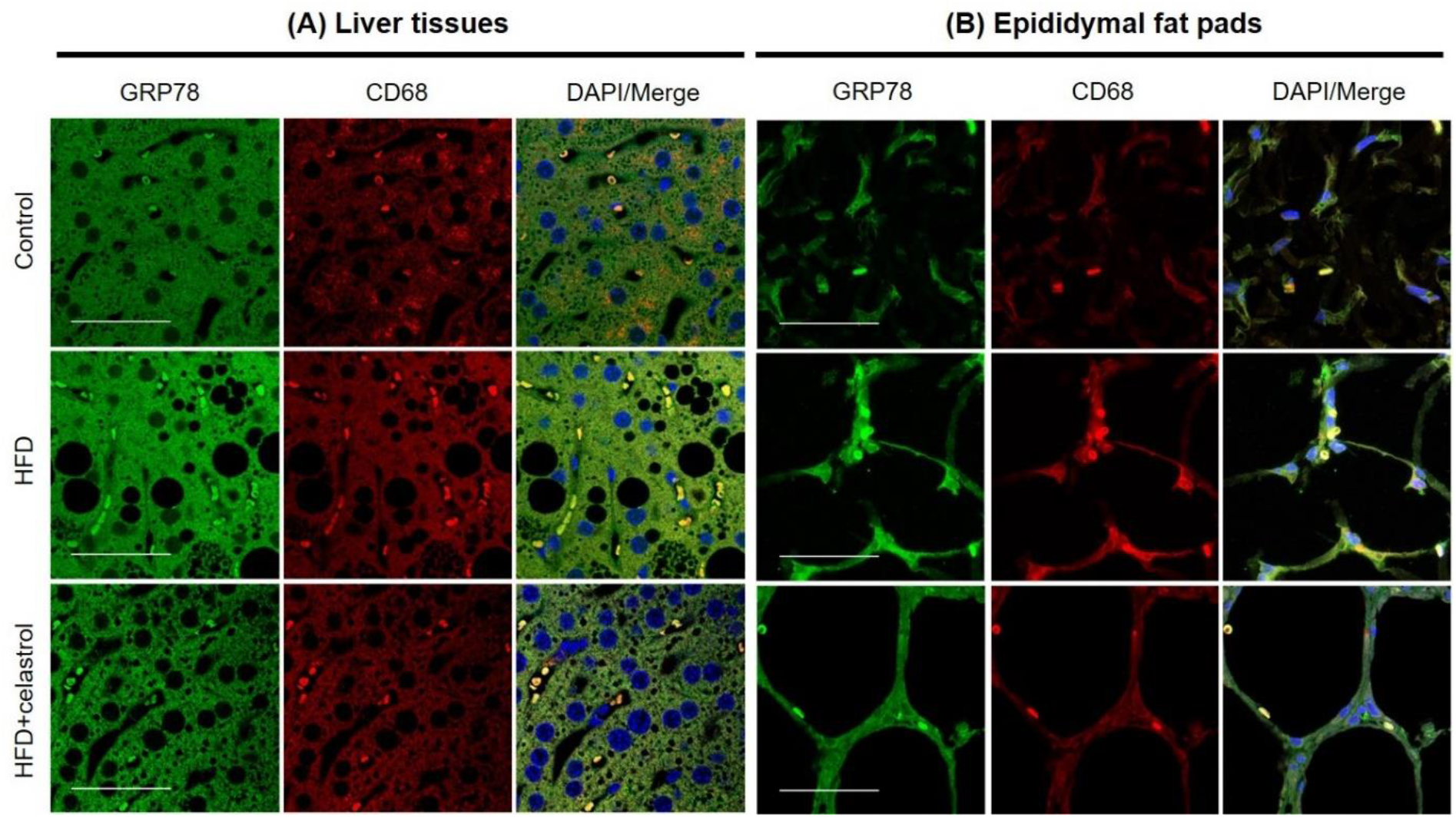
Celastrol reduces the expression of GRP78 in macrophages of liver and epididymal adipose tissues. (A) Detection of GRP78 in hepatic macrophages. After 21-day treatment, mouse livers were recovered and stained with antibodies against GRP78 and CD68, whereas DAPI was used to stain cell nuclei. The sections were imaged under a Zeiss LSM 780 confocal microscope. Representative images were shown. Scale bar, 50 μm. (B) Detection of GRP78 in adipose tissue macrophages. After 21-day treatment, mouse epididymal fat pads were recovered and stained with antibodies against GRP78 and CD68, whereas DAPI was used to stain cell nuclei. The sections were imaged under a Zeiss LSM 780 confocal microscope. Representative images were shown. Scale bar, 50 μm. **Source data 1.** Data for figure 3 A. **Source data 2.** Data for figure 3 B.

### RNA-Seq transcriptome profiling confirms the effects of celastrol on inflammation, ER stress and lipid accumulation in liver

To discover the effects of celastrol on hepatic inflammation, ER stress and lipid metabolism, we determined the effects of Nano-celastrol on the transcriptomic profile in mouse liver by next generation RNA-Seq technology. Based on the expression values and fold of change, a total of 49 differentially expressed genes (DEGs) were initially identified in the livers from the untreated control, HFD and HFD + Nano-celastrol groups. As shown in Figure 4A, 49 DEGs were compared by hierarchical clustering while the effects of HFD and celastrol on gene expression were presented in a heatmap. HFD up-regulated nine DEGs including Apoa4, Apoc2, Thrsp while Nano-celastrol suppressed the up-regulation of these nine genes and even down-regulated three genes (i.e., Bhmt, Car3, Mup18) in HFD-treated mice. By contrast, HFD down-regulated twenty-one DEGs (e.g., Cyp4a14, Mup7, Mup14, Mup15) while Nano-celastrol increased the expression of the selected genes to a certain extent and even up-regulated four genes (i.e., Hamp, Mt2, Igfbp2, Serpina1e) in HFD-treated mice. Among the other nineteen genes, Nano-celastrol significantly up-regulated four DEGs (i.e., Lrg1, Fgl1, Mt1, Apom) and down-regulated Scd1 although HFD showed little effect on these genes. Secondly, functional enrichment analysis of DEGs was performed by functional annotation tool DAVID bioinformatics resource 6.8 (https://david.ncifcrf.gov/). As shown in Figure 4B, Nano-celastrol significantly affected six KEGG pathways,14 biological processes and 5 molecular function groups. Interestingly, these functional groups mainly deal with retinal metabolism, lipid metabolism, fatty acid oxidation, cholesterol efflux, glucose metabolism, stress response, inflammatory response, oxidative stress, detoxification, PPAR pathway, kinase pathways, complement and coagulation cascade. Thirdly, these DEGs were analyzed for protein-protein interaction networks by online STRING tool (https://string-db.org/). As shown in Figure 4C, Nano-celastrol considerably affected four networks of gene products including ten DEGs for Network-1 (cyp1a2, cyp2a12, cyp3a25, cyp4a10, cyp4a14, cyp2c44, mdh1, pck1, ugt2b5); eight DEGs for Network-2 (apoa4, apoc2, apom, c3, c8a, creg1, itih3, itih4); three DEGs for Network-3 (acat1, ehhadh, hsd17b10) and two DEGs for Network-4 (mt1, mt2).

**Figure 4.**
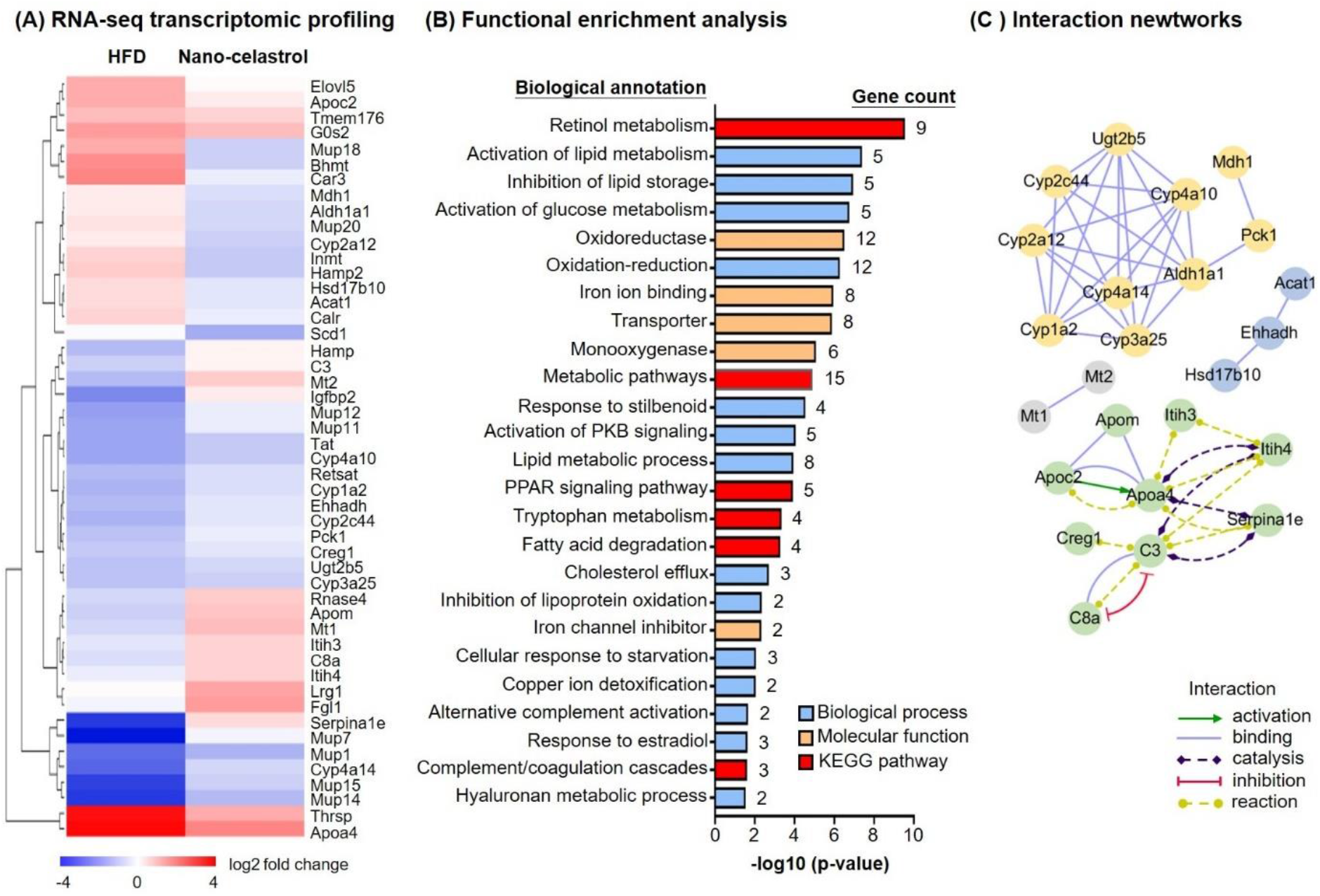
RNA-seq transcriptome profiling of celastrol-regulated genes. (A) Hierarchical cluster analysis of celastrol-regulated DEGs in HFD-induced obese mice. Expression values are determined by next generation RNA-seq technology on Illumina HiSeq2500 platform (www.genomics.cn). Data was presented as the expression values (log2 fold change) of the genes in HDF group and HFD+Nano-celastrol group relative to that in Control group. Increases in log2 fold of change are indicated in red hues while decreases are in blue hues. (B) Functional enrichment analysis of celastrol-regulated DEGs. Celastrol-regulated DEGs were classified into different functional groups in terms of molecular function, biological process and KEGG pathway using online DAVID v6.8. The number of genes related to each term were shown next to the bar. (C) Protein-protein interaction networks of the DEGs. Protein-protein interactions were analyzed by online STRING tool. The interaction networks involving DEGs were represented in different colors and lines using Cytoscape. **Source data 1.** Data for figure 4 A-C. **Source data 2.** Differentially expressed genes list for comparison.

### Celastrol ameliorated ER stress in palmitate-challenged macrophages

To verify the effects of celastrol on ER stress, we pretreated RAW264.7 macrophages with celastrol at the concentrations of 0.25, 0.5, 0.75 and 1 μM for 8 h and incubated with fatty acid palmitate (400 μM) for another 16 h to induce ER stress. Subsequently, the cellular proteins were extracted and analyzed by Western blotting for the expression of ER stress biomarkers (i.e., GRP78, IRE1α, ATF6, eIF2α, phospho-eIF2α, CHOP and XBP1). As shown in Figure 5A and 5B, plamitate not only elevated the expressions of ER stress biomarkers (i.e., GRP78, IRE1α, ATF6, CHOP and XBP1) but also induced the phosphorylation of eIF2α. Interestingly, celastrol effectively inhibited the expressions of GRP78, IRE1α, ATF6, CHOP and XBP1, and attenuated the phosphorylation of eIF2α in a concentration-dependent manner.

**Figure 5.**
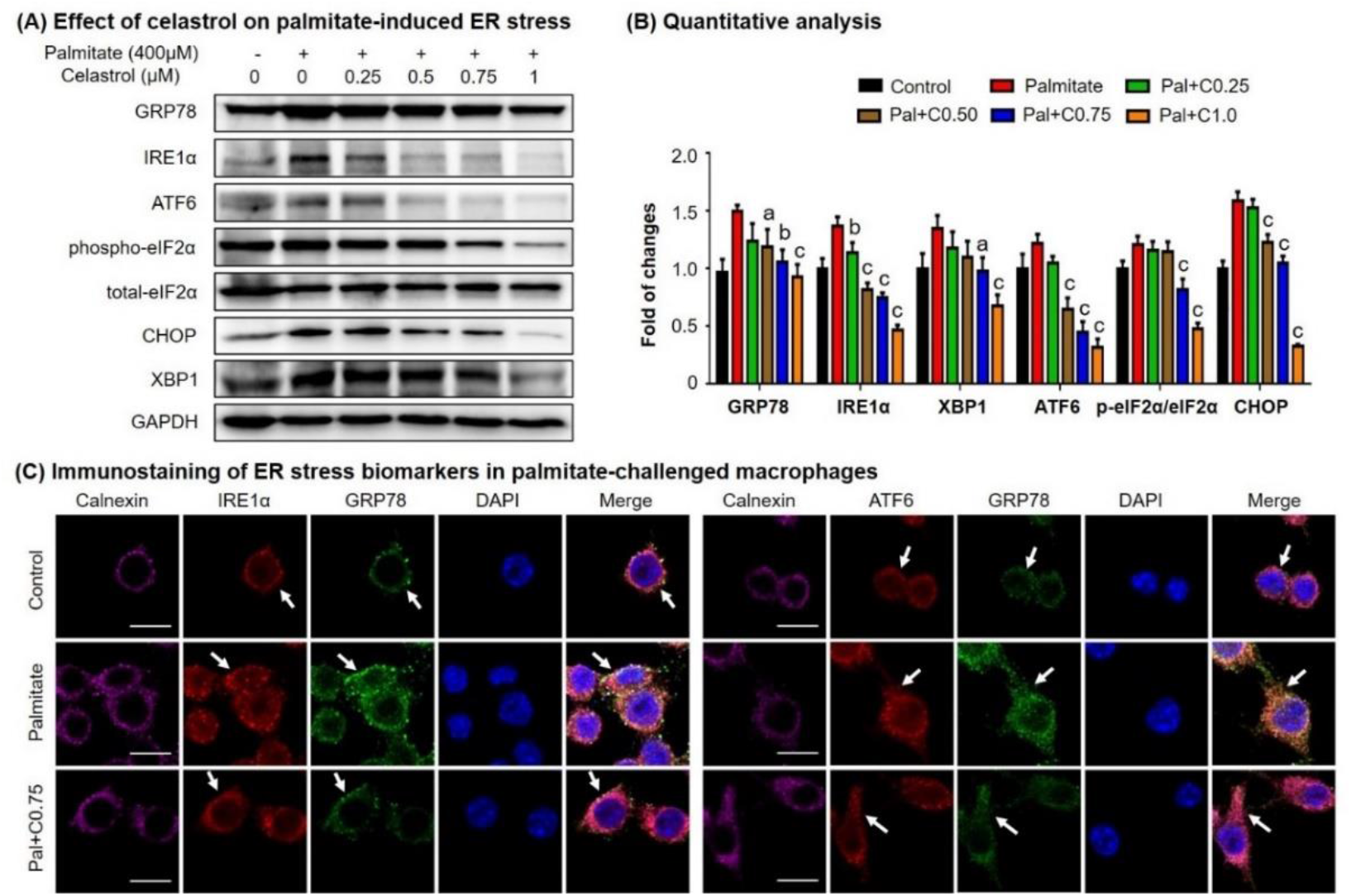
**Celastrol attenuates palmitate-induced ER stress in RAW264.7 macrophages.** (A) Western blot analysis for the effects of celastrol on the ER stress biomarkers in palmitate-challenged RAW264.7 cells. RAW264.7 cells were treated with celastrol for 8 h and subsequently with palmitate for another 16 h. The cellular proteins were analyzed by Western blotting with specific antibodies. Representative blots were shown. (B) Quantitative analysis for the expression levels of ER stress biomarkers. The blots (n = 3) were quantified by a densitometric method. C0.25, celastrol (0.25 μM); C0.5, celastrol (0.5 μM); C0.75, celastrol (0.75 μM); C1.0, celastrol (1 μM); a, *p*<0.05; b, *p*<0.01; c, *p*<0.001. (C) Co-localization of GRP78, IRE1α and ATF6 in palmitate-challenged RAW264.7 cells. After the pretreatment with celastrol for 8 h and subsequently with palmitate for another 16 h, the cells were stained with antibodies against GRP78, IRE1α, ATF6 and calnexin, whereas DAPI was used to stain the cell nuclei. The cells were imaged under a Zeiss LSM 780 confocal microscope. Representative images were shown. C0.75, celastrol (0.75 μM); Palmitate (400 μM). Scale bar, 10 μm. **Source data 1.** Original immunoblots. **Source data 2.** Data for Figure 5B.

To validate the inhibitory effects of celastrol on ER stress, we detected the cellular levels of GRP78, IRE1α, ATF6 and calnexin by immunofluorescence staining. Following the pretreatment with 0.75 μM celastrol for 8 h and co-incubation with palmitate for another 16 h, RAW264.7 cells were stained with antibodies against GRP78, IRE1α, ATF6, whereas calnexin was detected as a membrane marker of ER. As shown in Figure 5C, palmitate significantly increased the expression of GRP78, IRE1α, ATF6, while celastrol resored the expression of these proteins to the control levels. Moreover, palmitate caused the dissociation of GRP78 from ER membrane senors IRE1 and ATF6, whereas celastrol maintained the colocalization of GRP78 with IRE1 and ATF6 in ER compartment.

### Identification of celasrol-GRP78 conjugate

To discover the molecular targets of celastrol, we synthesized a new celastrol-PEG4-alkyne bearing an alkyne (-C≡C-) group as a small molecular probe. As shown in Figure 6A, following drug treatment, the celastrol-bound proteins were biotinylated through Click chemistry “azide-alkyne cycloaddition”, and separated by affinity enrichment with streptavidin-coated magnetic beads. As shown in Figure 6B, ER chaperone GRP78 was identified as a major celastrol-bound protein by mass spectrometry (MS) and verified by Western blotting with anti-GRP78 antibody. The binding of celastrol to GRP78 was characterized by peptide mapping and virtual simulation with AutoDock Vina in PyRx-virtual screen tool package. For peptide mapping, celastrol was incubated with recombinant mouse GRP78, digested with trypsin and analyzed by LC-MS/MS technology. As shown in Figure 6C, celastrol covalently bound to Cys^41^ in the peptide with the sequence of GSEEDKKEDVGTVVGIDLGTTYSC^41^VGVFK. As shown in Figure 6D, molecular docking also supports that celastrol binds to the Cys^41^-containing binding site in GRP78 structure with the binding energy of −8.1 kcal·mol^-1^ and the inhibition constant of 1.55 μM.

**Figure 6.**
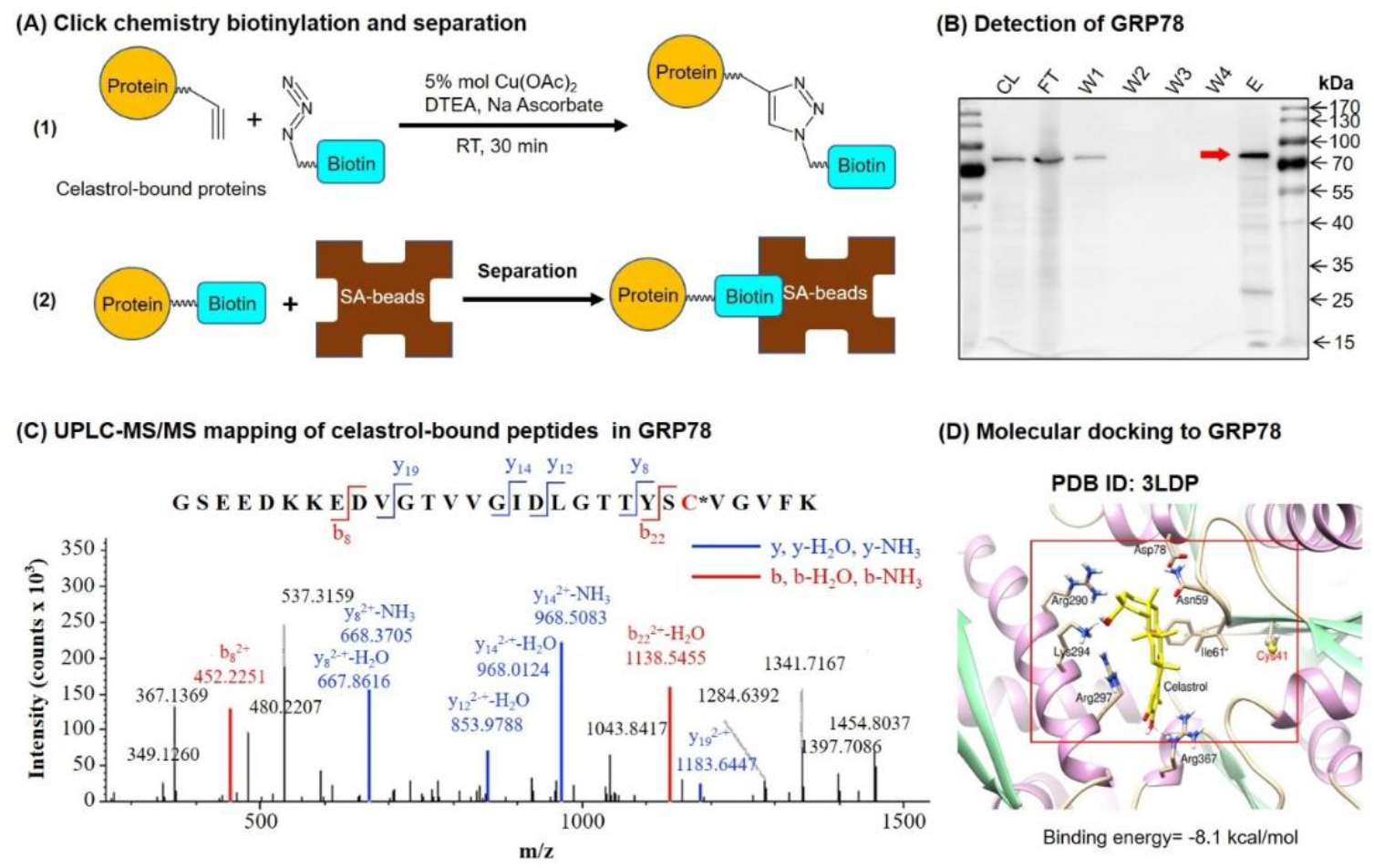
Celastrol forms covalent conjugate with ER chaperone GRP78. (A) Click chemistry biotinylation and affinity isolation of celastrol-bound proteins. Following 24 h treatment with celastrol-PEG4-alkyne, the cellular proteins were isolated from RAW264.7 cells and biotinylated with Azide-PEG3-Biotin under Click chemistry conditions. The biotinylated proteins were purified by binding to streptavidin-coated magnetic beads. (B) Western blot verification of celastrol-bound proteins. After Click chemistry biotinylation and affinity isolation, celastrol-bound proteins were resolved by 10% SDS-PAGE and visualized by immunoblotting with anti-GRP78 antibody. (C) Mapping of celastrol-binding site in GRP78. Recombinant mouse GRP78 was prepared from DE3 *E. coli* cells and incubated with celastrol in 50 mM NaHCO_3_ buffer containing 10% DMSO. After resolution by Clear-Native-PAGE and Coomassie blue staining, celastrol-GRP78 conjugate was digested with trypsin and analyzed by LC/MS/MS technology. (D) Molecular simulation of celastrol-GRP78 interactions. Celastrol was docked into the crystal structure of GRP78 (PDB ID: 3LDP) using Autodock vina in PyRx 0.8 (http://pyrx.sourceforge.net/downloads). **Source data 1.** Original immunoblots and figure with the uncropped blots with the relevant bands labelled. **Source data 2.** Data for Figure 6C.

### Celastrol attentuated the binding of peptide to GRP78

To clarify the effects of celastrol on the chaperone activity of GRP78, we employed surface plasmon resonance (SPR) binding analysis technology to determine the binding of synthetic GRP78-targeting peptide “WDLAWMFRLPVG” to GRP78 on Biacore X100 SPR analyzer. The affinity was determined by steady-state analysis while Kd values were estimated using Biacore X100 Evaluation Software. Biotinylated GRP78 was initially immobilized onto streptavidin-coated Sensor Chip SA, and synthetic peptide at a wide range of concentrations was subsequently pumped into the system. As shown in Figure 7A, the synthetic peptide binds to GRP78 with Kd value of 17.7 nM. As shown in Figure 7B, in contrast, when GRP78-coated Sensor Chip was saturated by 10 μM celastrol prior to the binding of GRP78-targeting peptide, the peptide exhibited Kd value of 2.23 μM.

**Figure 7.**
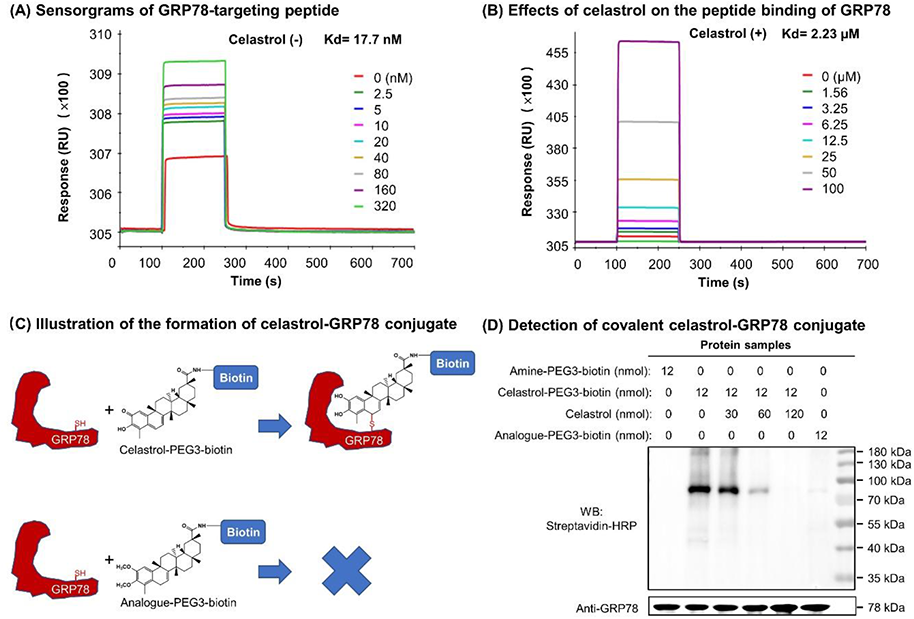
Covalent binding of celastrol attenuated the chaperone activity of GRP78 towards peptide. (A) Validation of peptide-binding to GRP78. After the immobilization of biotinylated GRP78 to the streptavidin-coated Sensor Chip SA, GRP78-targeting peptide (WDLAWMFRLPVG) was pumped into the Biacore X100 SPR system for evaluating the peptide binding to GRP78 under steady-state condition. After four runs, the average Kd value was calculated by Biacore X100 Evaluation Software. Representative sensorgram was shown. (B) Effects of celastrol on peptide binding to GRP78. After the saturation with 10 μM celastrol, GRP78-targeting peptide (WDLAWMFRLPVG) (n=4) was pumped into the Biacore X100 SPR system for calculating the average Kd value for peptide binding to GRP78. Representative sensorgram was shown. (C) Scheme illustrating the formation of covalent celastrol-GRP78 conjugate. The cysteine of GRP78 reacts with quinone methide moeity of celastrol whereas the analogue lacking quinone methide structure fails to form covalent conjugate with GRP78. (D) Detection of covalent celastrol-GRP78 conjugate. Following the incubation of recombinant GRP78 with celastrol-PEG3-biotin or analogue-PEG3-biotin, the reaction mixtures were analyzed by Western blotting with streptavidin-HRP or anti-GRP78 antibody. **Source data 1.** Data for Figure 7 A-B. **Source data 2.** Original immunoblots and figure with the uncropped blots with the relevant bands labelled.

### Quinone methide is required to form covalent celastrol-GRP78 conjugate

To determine the involvement of quinone methide in covalent conjugation, celastrol was reduced by NaBH_4_ and protected by O-methylation to yield celastrol analogue (Figure 7C), which lacks the chemically reactive quinone methide moiety. When recombinant GRP78 was incubated with excessive celastrol-PEG3-biotin, celastrol analogue-PEG3-biotin or amine-PEG3-biotin, celastrol-PEG3-biotin was readily attached to GRP78 whereas neither of celastrol analogue-PEG3-biotin or amine-PEG3-biotin was detectable in Western blot analysis (Figure 7D). Free celastrol effectively competed with celastrol-PEG3-biotin for covalent conjugation in a concentration-dependent manner while biotinylation was completely blocked by celastrol at 10-fold excess of molar concentration. On the other hand, the covalent celastrol-GRP78 conjugates in RAW264.7 macrophages were visualized by labeling with a highly reactive probe AFDye555-picolyl azide under Click chemistry conditions as described (Uttamapinant et al. 2012). Upon the formation of covalent celastrol-GRP78 conjugate, celastrol-PEG4-alkyne could react with AFDye555-picolyl azide through “Alkyne-Azide cycloadduction”. As shown in Figure 8, the celastrol-PEG4-alkyne-treated cells indeed showed strong red fluorescence whereas the celastrol-treated cells did not show any red fluorescence. Importantly, the labeling of celastrol-PEG4-alkyne appeared to be well colocalized with GRP78 and Calnexin.

**Figure 8.**
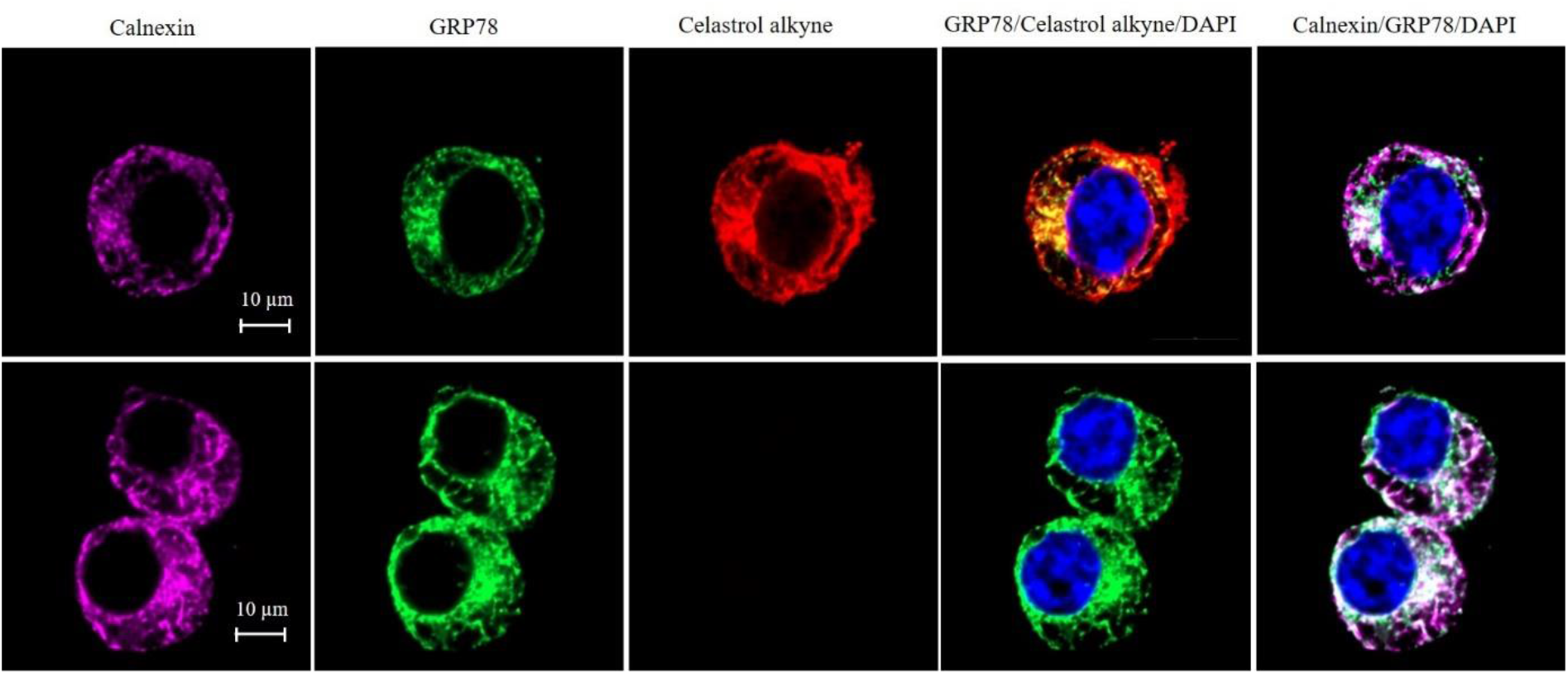
Intracellular colocalization of celastrol labeling and GRP78. Following the challenge with palmitate, RAW264.7 cells were treated with celastrol or celastrol-PEG4-alkyne, washed, fixed, reacted with fluorescence dye AFDye555-picolyl azide. The cells were immunostained with antibodies against GRP78 and calnexin, fluorescent secondary antibodies whereas DAPI was used to stain the cell nuclei. The cells were imaged under a Zeiss LSM 780 confocal microscope. Representative images were shown. Scale bar, 10 μm. **Source data 1.** Original immunofluorescence staining result of RAW264.7 cells treated with celastrol-PEG4-alkyne. **Source data 2.** Original immunofluorescence staining result of RAW264.7 cells treated with celastrol.

### Quinone methide is required to induce weight loss in diet-induced obese mice

To clarify the importance of covalent GRP78 inhibition in ani-obesity effects, the diet-induced obese mice were treated with either celastrol or celastrol analogue at the dose of 5 mg/kg/day via oral gavage for consecutive 21 days. Based on the measurement of body and body fat contents as shown in Figure 9A-9D, celastrol effectively reduced the body weight and fat mass while recovered the loss of lean mass in HFD-induced obesity. In contrast, celastrol analogue lacking quinone methide moiety failed to ameliorate weight gain, fat accumulation and lean mass loss. Furthermore, the ipGTT and ITT results in Figure 9E&9F showed that celastrol analogue could no longer ameliorate glucose tolerance and insulin tolerance compared with celastrol.

**Figure 9.**
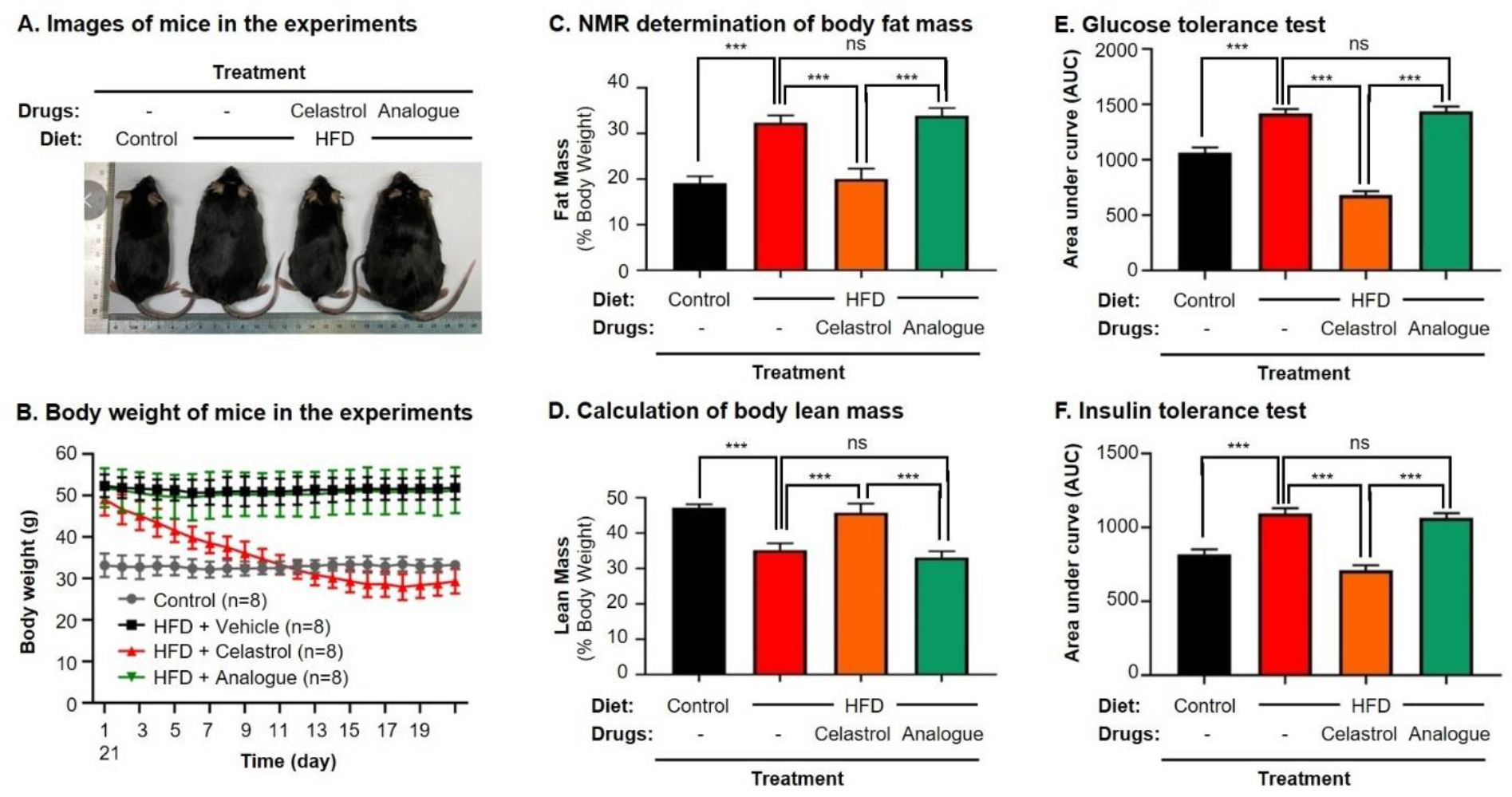
Quinone methide is essential to induce weight loss, reduce fat mass and restore glucose tolerance and insulin tolerance in diet-induced obese C57BL/6N mice. (A) Images of experimental mice. Diet-induced obese mice were treated with celastrol at the doses of 0, 5.0, 7.5 mg/kg/d for 21 days, whereas control mice were fed with normal diet. Experimental mice were imaged at day-21. (B) Measurement of body weight. Body weight was daily measured (n=8). (C) NMR determination of fat contents. After 21-day treatment, fat contents in living mice were analyzed by Bruker minispec NMR analyzer. (D) Calculation of body lean mass. Lean mass was calculated from NMR determination of body fat mass as described in (C). Body weight was daily measured. (E) Glucose tolerance test (GTT). After glucose injection, blood glucose levels were measured and plotted against time. The area under curve (AUC) for each group was calculated. (F) Insulin tolerance test (ITT). After insulin injection, blood glucose levels were measured and plotted against time. The AUC for each group was calculated. ***, *p* < 0.001; ns, not significant. **Source data.** Data for Figures 9B-F.

## Discussion

Pharmacological modulation of ER stress is a potential therapeutic strategy to treat various metabolic disorders (Cao and Kaufman 2013). Chaperone protein GRP78 is essential to regulate ER stress, inflammation and lipid metabolism since genetic knockout of GRP78 abolishes the growth, differentiation and maturation of preadipocytes (Zhu et al. 2013). By contrast, several small molecule GRP78 inhibitors were recently developed for restoring ER homeostasis, especially, KP1339/IT-139 was evaluated for anticancer activity in a phase I clinical trial (Schoenhacker-Alte et al. 2017, Chen et al. 2018, Gurusinghe et al. 2018). Other purported GRP78 inhibitors such as Verrucocidin also ameliorate ER stress through different mechanisms other than direct binding to GRP78 (Thomas et al. 2013). Currently, little is known about the potential of covalent GRP78 inhibition in the regulation of ER stress. Here we determined whether celastrol could ameliorate ER stress, inflammation and metabolic dysfunctions in diet-induced obesity via covalent modification of chaperone GRP78.

HFD is well-known to induce the deposition of lipids and alter the composition of fatty acids in liver, adipose tissues and other organs (Cameron-Smith et al. 2003, Oosterveer et al. 2009, Zhao et al. 2019). Based on transcriptomic profiling and biochemical analysis, HFD up-regulates several lipogenic genes for producing fatty acids (Acc, Fas, Scd1, Elovl6) and triglycerides (Dgat, Gpat), elongase for synthesizing long chain fatty acids such as n-3 and n-6 polyunsaturated fatty acids, and Srebp-1c and Pgc-1β for lipid uptake (Oosterveer et al. 2009). Free fatty acids induce ER stress and inflammatory response in macrophages and adipose tissues via TLR-mediated mechanisms (Han and Kaufman 2016). ER stress in turn orchestrates unfolded protein response, inflammation, metabolic dysfunctions and insulin resistance in obesity and diabetes (Ozcan et al. 2004, Hummasti and Hotamisligil 2010). Our group and others have recently demonstrated that celastrol effectively reduces adipose hypertrophy in mouse and rats models of HFD-induced obesity (Wang et al. 2014, Zhang et al. 2017, Zhao et al. 2019). In the present study, firstly, H&E staining confirmed that celastrol ameliorated adipose hypertrophy and reduced lipid accumulation in liver and epididymal adipose tissues (Figure 1A and 1B). Secondly, GC-MS technology was employed to profile fatty acids in liver and adipose tissues from Control, HFD and HFD + Celastrol groups. Based on the quantitative analysis in Figure 1E and 1F, celastrol ameliorated HFD-induced elevations of various polyunsaturated fatty acids (e.g., C16:1, trans-C18:1, cis-C18:1, C18:2, C18:3, cis-C20:2, cis-C20:3) in liver but showed little activity against HFD-impaired production of long chain polyunsaturated fatty acids eicosoids (e.g., cis-C20:1, cis-C20:3, cis-C22:6) in epididymal fat pads. These results suggest that celastrol may mainly act on the metabolism of fatty acids in liver. Thirdly, qRT-PCR technique was employed to determine the key biomarkers for adipogensis, hormone secretion and energy expenditure. The key findings include: 1) celastrol could effectively antagonize HFD-induced upregulation of adipogenic genes (e.g., PPAR-γ, C/EBP-α, aP2) and adipocyte-secreted hormones (e.g., leptin, resistin, adipisin) in livers shortly after 7-day treatment; 2) celastrol markedly elevated the expression of lipolytic genes (e.g., CTP-1, CAT, ACO) and thermogenic gene UCP2 in a dose-dependent manner (Figure 2A and 2B); 3) celastrol mainly suppressed HFD-induced upregulation of adipogenic genes (e.g., PPAR-γ, C/EBP-α, aP2) and adipocyte-secreted hormones (e.g., leptin, resistin, adipisin) in epididymal fat pads (Figure 2C). These results suggest that celastrol mainly reprograms the biosynthesis and degradation of fatty acids in liver while inhibits fatty acid biosynthesis and hormone secretion. On the other hand, the activation of ER stress sensor IRE1α exacebates obesity-associated inflammation (Shan et al. 2017). However, celastrol could reduce ER stress via reducing the protein level of p-PERK/PERK in hypothalamus of diet-induced obese mice (Liu et al. 2015). Along this line, we examined the *in vivo* effects of celastrol on ER stress by detecting the expression of macrophage biomarker CD68 and ER stress biomarker GRP78 in hepatic and adipose tissues from the obese mice. HFD markedly upregulated the expression of GRP78 in hepatic and adipose tissue macrophages, while celastrol at the dose of 7.5 mg·kg^-1^·d^-1^ restored the expression of GRP78 in macrophages (Figure 3). These results indicate that celastrol could ameliorate HFD-induced ER stress in hepatic and adipose tissue macrophages. To facilitate safe control of body weight, we recently prepared non-toxic Nano-celastrol nanoparticles by entrapping celastrol in PEG-PCL nanomicelles for the treatment of diet-induced obesity (Zhao et al. 2019). In the present study, we further identified 49 DEGs in the livers from mice in Control, HFD and HFD+Nano-celastrol groups by next generation RNA sequencing technology (Figure 4A). Indeed, Figure 4B clearly shows that these DEGs are mainly involved in the regulation of retinol metabolism (i.e., Aldh1a1, Cyp4a10, Cyp1a2, Cyp2a12, Cyp2c44, Cyp3a25, Cyp4a14, Retsat, Ugt2b5), lipid biosynthesis (e.g., Apoc2, C3, Cyp1a2, Cyp4a10, Cyp4a14, Ehhadh, Elovl5, Pck1, Scd1, Thrsp), oxidative stress (e.g., Aldh1a1, Creg1, Cyp1a2, Cyp2a12, Cyp3a25, Cyp4a10, Cyp4a14, Ehhadh, Hsd17b10, Mdh1, Retsat, Scd1), lipid degradation (i.e., Acat1, Cyp4a10, Cyp4a14, Ehhadh), glucose metabolism (i.e., Mup1, Mup11, Mup12, Mup18, Mup20), cholesterol efflux (i.e., Apoa4, Apoc2, Apom) and inflammatory response (e.g., C3, C8a, Serpina1e) although some DEGs also contribute to several other biological processes and pathways. By using online software STRING, functional enrichment analysis (Figure 4C) reveals four protein-protein interaction networks: 1) lipid and glucose metabolism (i.e., Mdh1, Pck1, Aldh1a1, Cyp1a2, Cyp2a12, Cyp2c44, Cyp3a25, Cyp4a10, Cyp4a14, Ugt2b5); 2) apolipoprotein metabolism and inflammatory response (i.e., Apoa4, Apoc2, Apom, Ltih3, Ltih4, Serpina1e, C3, Creg1, C8a); 3) mitochondrial energy homeostasis (i.e., Acat1, Enhadh, Hsd17b10); 4) adipose tissue energy homeostasis (Mt1, Mt2). Thus, RNA-Seq profiling essentially supports the potential of celastrol in the regulation of ER stress, lipid accumulation and inflammatory response in the liver.

ER chaperone protein GRP78 is integral to the celluluar sensing of fatty acid-induced ER stress and inflammatory stimuli and the regulation of lipid uptake, adipogenesis, lipolysis, glucose metabolism, cholesterol efflux, inflammatory response and immune response (Zhu et al. 2013). Moreover, GRP78 and CHOP modulate macrophage apoptosis and control the progression of bleomycin-induced pulmonary fibrosis (Ayaub et al. 2016). In the present study, RAW264.7 macrophages were treated by fatty acid palmitate to induce ER stress. Following celastrol treatment, we subsequently employed Western blotting and fluorescence immunostaining to determine the expression levels of ER stress biomarkers. As result, celastrol not only reduced the levels of GRP78, IRE1α, ATF6, CHOP and XBP1, but also inhibited palmitate-stimulated eIF2α phosphorylation in a concentration dependent manner (Figure 5A and 5B). Based on the immunostaining results (Figure 5C), celastrol decreased the signal intensity of GRP78, ATF6 and IRE1α and preserved the co-localization of GRP78 with ATF6 and IRE1α in the ER compartment. These results indicate that celastrol could effectively ameliorate ER stress in hepatic and adipose tissue macrophages.

Celastrol bears a chemically reactive quinone methide moeity and thereby tends to form covalent conjugate with certain proteins such as HIV-Tat, IKKalpha and beta kinases, Cdc37 and HSF although such covalent attachment appears to be reversible (Salminen et al. 2010, Narayan et al. 2011, Zhang et al. 2018). However, it is critical to identify protein targets with convincing association to ER signaling, anti-inflammatory and metabolic reprogramming. In the present study, a novel celastrol-PEG4-alkyne bearing an alkyne (-C≡C-) group was firstly synthesized as a small molecule probe through the in-house synthetic procedure (Cheng et al. 2020). Secondly, RAW264.7 macrophages were treated with celastrol-PEG4-alkyne and lysed. The cellular proteins were biotinylated via Click chemistry “azide-alkyne cycloaddition” and separated by affinity enrichment onto streptavidin-coated magnetic beads (Figure 6A). The protein fractions were resolved by 10% SDS-PAGE and identified by MALDI-MS technology. The major protein band was identified to be GRP78 and subsequently verified by Western blotting using specific antibody (Figure 6B). Thirdly, recombinant mouse GRP78 protein was prepared in *E. coli* BL21 cells and incubated with celastrol. Following trypsin digestion, peptides were mapped by LC-MS/MS analysis. Peptide mapping revealed a celstrol-bound peptide with the sequence of GSEEDKKEDVGTVVGIDLGTTYSC^41^VGVFK (Figure 6C). This experiment suggests that celastrol interacts with GRP78 by covalent attachment to Cys^41^. Forthly, the binding of celastrol to GRP78 was simulated by molecular docking. Interestingly, celastrol binds to the Cys^41^-containing region in GRP78 structure with binding energy of −8.1 kcal/mol and inhibition constant of 1.55 μM (Figure 6D). Fifthly, a recently identified peptide WDLAWMFRLPVG was synthesized to examine the effects of celstrol on the chaperone activity of GRP78 as described (Arap et al. 2004). Biacore X100 SPR system was employed to evaluate whether celastrol could alter the binding of peptide WDLAWMFRLPVG to GRP78 on the Sensor chip. Indeed, celastrol attenuated the chaperone activity of GRP78 since the average Kd value of peptide binding was increased from 17.7 nM to 2.23 μM (Figure 7A and 7B). Sixthly, two new celastrol derivatives were prepared to explore the importance of quinone methide moiety for celastrol to bind to GRP78 (Figure 7C). Celastrol-PEG3-biotin was readily detected in association with GRP78 although the binding could be blocked by excess celastrol. By contrast, analogue-PEG3-biotin failed to bind to GRP78 due to the lack of quinone methide moiety (Figure 7D). Seventhly, the present study addressed whether the formation of celastrol-GRP78 conjugate could be colocalized with the intracellular GRP78 (Figure 8). The cells were treated with celastrol-PEG4-alkyne and subsequently labeled with fluorescence dye AFDye555-picolyl azide whereas GRP78 and another ER biomarker Calnexin were detected by immunostaining. As result, the fluorescence of celastrol-PEG4-alkyne labeling was well colocalized with GRP78, and, to a large extent, ER biomarker Calnexin. Last but not least, the present study determined whether quinone methide moirty was essential for anti-obesity effects (Figure 9). Celastrol was reduced by NaBH_4_ and protected by O-methylation to yield celastrol analogue. Unlike celastrol, such analogue became totally inactive to induce weight loss, reduce fat mass and ameliorate glucose tolerance and insulin tolorance in diet-induced obese mice as assessed by previous described procedure (Luo et al. 2017, Zhao et al. 2019).

In summary, the present study demonstrated that celastrol ameliorated adipose hypertrophy and reduced lipid accumulation in liver and adipose tissues of diet-induced obese mice. Celastrol might induce rapid reduction of body weight through suppressing lipid uptake, lipogenesis and increasing lipolysis and thermogenesis. The key finding was that celastrol formed covalent conjugate with chaperone protein GRP78 in macrophages, diminished the chaperone activity of GRP78 and reduced the activation of ER stress sensors against lipidotoxicity. The present study suggests that the covalent inhibition of GRP78 is a novel anti-obesity mechanism for reprograming the signaling networks that control inflammation, lipid accumulation, ER stress in liver and adipose tissues. Ultimately, celastrol becomes an important lead compound for the development of new selective, irreversible GRP78 inhibitors to treat obesity, diabetes and related cardiovascular diseases.

## Methods

### Antibodies and biochemical reagents

Antibodies against GRP78, IRE1α, XBP1 and calnexin were purchased from Abcam (Cambridge, UK). Antibodies against ATF6, CHOP, phospho-eIF2α, eIF2α, XBP1 and GAPDH were purchased from Cell Signaling Technology (Boston, MA, USA). Alexa Fluor 594 and 568-conjugated goat anti-rabbit IgG secondary antibodies and Alexa Fluor 488-conjugated goat anti-mouse IgG secondary antibody were purchased from Invitrogen (Carlsbad, CA, USA), while Alexa Fluor 647-conjugated donkey anti-goat IgG secondary antibody was purchased from Abcam (Cambridge, UK). Dulbecco’s modified Eagle’s Medium (DMEM), fetal bovine serum (FBS) and 100x penicillin and streptomycin solution were purchased from Invitrogen (Carlsbad, CA, USA). Protein Assay Dye Reagent Concentrate was purchased from Bio-Rad (Hercules, CA, USA). Celastrol with the purity of over 98% (HPLC) was purchased from Nanjing Spring and Autumn Biological Engineering Co., Ltd. (Nanjing, China). Anti-rabbit HRP-conjugated IgG secondary antibody, bovine serum albumin (BSA), insulin, lipopolysaccharides (LPS), dexamethasone, indomethacin, sodium palmitate and other chemicals were purchased from Sigma-Aldrich (St. Louis, MO, USA) unless indicated otherwise.

### Chemical synthesis of celastrol derivatives

Celastrol was purchased from Nanjing Spring & Autumn Biological Engineering Company (Nanjing, Jiangsu, China). Biotin-PEG3-amine and Alkyne-PEG4-amine were purchased from Hunan Huateng Pharmaceutical Company (Changsha, Hunan, China). The solvents and reagents including 4-methoxybenzyl chloride (PMB-Cl), iodomethane (CH_3_I), (benzotriazol-1-yloxy)tris(dimethylamino)phosphonium hexafluorophosphate (BOP), sodium bicarbonate (NaHCO_3_), potassium carbonate (K_2_CO_3_), sodium borohydride (NaBH_4_), dimethylformamide (DMF), methanol (MeOH), chloroform, dichloromethane (DCM), trifluoroacetic acid (TFA) and triethylamine (TEA) were purchased from Sigma Aldrich (St Louis, MO, USA). The solvents for extraction and silica gel column chromatography were purchased from RCI Labscan Limited (Bangkok, Thailand) or Duksan Pure Chemicals Company (Ansan, South Korea). All solvents and reagents were used without further preparation unless otherwise indicated. The NMR ^1^H and ^13^C spectra were recorded on Bruker AVANCE III HD 400 MHz spectrometer. The chemical shifts were reported in ppm relative to the internal standard signal. The peak splitting patterns were described as follows: s = singlet, d = doublet, t = triplet, q = quartet, m = multiplet, while the coupling constant, *J*, was reported in Hertz unit (Hz). The molecular mass was determined on AB SCIEX X500R Q-TOF mass spectrometer in positive ion mode or negative ion mode. The purity of the products was determined by HPLC analysis on a reversed phase ACE Excel C18 column (2 μm particle size, 100 × 2.1 mm, Advanced Chromatography Technologies Ltd., Scotland) under the control of UltiMate 3000 UPLC system (Thermo Fisher Scientific, USA) equipped with quaternary pump, autosampler, thermostat column compartment and diode array detection system. The products were generally prepared at the purity of >=95% based on the integration of UV spectra at 254 nm.

Celastrol analogue (2,3-dimethoxydihydrocelastrol) was synthesized from celastrol through several steps of chemical reaction and purified by silica gel column chromatography in the solvents of PE/EA (6:1). Yield: 325 mg, 0.68 mmol, 81.9 %. ^1^H NMR (400 MHz, Chloroform-*d*) δ 6.65 (s, 1H), 5.64 (s, 1H), 3.71 (s, 3H), 3.63 (s, 3H), 3.15 (dd, *J* = 20.8, 6.0 Hz, 1H), 2.89 (d, *J* = 20.8 Hz, 1H), 2.32 (d, *J* = 15.4 Hz, 1H), 2.04 (s, 3H), 2.00 – 1.88 (m, 4H), 1.71 (ddd, *J* = 32.6, 16.0, 10.2 Hz, 2H), 1.57 – 1.46 (m, 2H), 1.48 – 1.37 (m, 3H), 1.30 (d, *J* = 15.9 Hz, 2H), 1.21 (s, 3H), 1.11 (s, 3H), 1.07 (s, 3H), 0.95 (s, 3H), 0.79 (d, *J* = 14.8 Hz, 1H), 0.62 (s, 3H). ^13^C NMR (101 MHz, CDCl_3_) δ 181.69, 150.62, 149.22, 144.89, 144.38, 127.63, 125.61, 117.47, 106.08, 60.20, 55.63, 44.24, 43.63, 40.09, 37.45, 37.10, 36.74, 34.67, 34.37, 33.87, 32.71, 31.39, 30.54, 30.41, 30.10, 29.71, 28.77, 27.72, 22.74, 18.32, 11.55. HRMS (m/z): [M-H]^-^ calcd. for C_31_H_43_O_4_, 479.3161; found, 479.3164.

**Supplementary figure source data 1.** Data for the structures, HRMS spectrum, ^1^H-NMR spectrum and ^13^C-NMR spectrum of celastrol analogue.

**Supplementary figure source data 2.** Synthesis route of celastrol analogue and data for the structures, HRMS spectrum, ^1^H-NMR spectrum, and ^13^C-NMR spectrum of intermediates.

Celastrol-PEG3-biotin (*N*-(2-(2-(2-((5-(2-oxohexahydro-1H-thieno[3,4]-dimidazol-4-yl) pentanamido) ethoxy) ethoxy) ethoxy) ethyl) celastrol-20-carboxamide) was prepared by direct conjugation of celastrol with biotin-PEG3-amine in dimethylformamide (DMF) in the presence of BOP and triethylamine as previously described (Cheng et al. 2020). The product was purified by silica gel column chromatography in the solvents of DCM/MeOH (50:1 to 10:1). Yield: 122 mg, 0.14 mmol, 60.5 %. ^1^H NMR (400 MHz, Chloroform-*d*) δ 7.94 – 7.58 (m, 1H), 7.46 – 7.28 (m, 1H), 7.01 (d, *J* = 6.6 Hz, 1H), 6.51 (dd, *J* = 12.3, 6.4 Hz, 2H), 6.32 (d, *J* = 7.2 Hz, 1H), 6.01 – 5.71 (m, 1H), 4.46 (dd, *J* = 8.0, 4.8 Hz, 1H), 4.27 (dd, *J* = 8.0, 4.7 Hz, 1H), 3.62 – 3.50 (m, 10H), 3.42 (d, *J* = 13.2 Hz, 6H), 3.33 – 3.25 (m, 2H), 3.09 (q, *J* = 7.3 Hz, 2H), 2.89 – 2.67 (m, 2H), 2.41 (d, *J* = 15.3 Hz, 1H), 2.19 (d, *J* = 6.2 Hz, 6H), 2.01 (t, *J* = 12.2 Hz, 2H), 1.89 – 1.77 (m, 3H), 1.67 – 1.60 (m, 4H), 1.55 (d, *J* = 7.6 Hz, 2H), 1.48 (q, *J* = 7.8, 6.2 Hz, 3H), 1.41 (s, 3H), 1.32 (t, *J* = 7.3 Hz, 2H), 1.23 (s, 3H), 1.12 (s, 3H), 1.08 (s, 3H), 0.60 (s, 3H). ^13^C NMR (101 MHz, CDCl_3_) δ 178.41, 178.29, 173.78, 170.80, 164.99, 164.23, 146.15, 134.54, 127.38, 119.55, 118.17, 117.58, 77.36, 70.43, 70.13, 69.74, 61.96, 60.35, 55.69, 50.72, 46.00, 45.18, 44.46, 43.15, 40.35, 39.46, 39.38, 39.23, 38.30, 36.45, 35.96, 35.06, 33.86, 33.64, 31.73, 31.19, 30.85, 30.09, 29.48, 28.75, 28.32, 28.15, 25.71, 21.80, 18.43, 10.40, 8.73. HRMS (m/z): [M+ H] ^+^calcd. for C_47_H_71_N_4_O_8_S, 851.4993; found, 851.3923.

**Supplementary figure source data 3.** Data for the structures, HRMS spectrum, ^1^H-NMR spectrum and ^13^C-NMR spectrum of celastrol-PEG3-biotin.

Analogue-PEG3-biotin (*N*-(2-(2-(2-((5-(2-oxohexahydro-1H-thieno[3,4]-dimidazol-4-yl)pentanamido)ethoxy)ethoxy)ethoxy)ethyl)2,3-dimethoxydihydrocelastrol-20-carboxamide) was prepared by direct conjugation of celastrol with biotin-PEG3-amine in DMF in the presence of BOP and TEA as previously described (Cheng et al. 2020). The product was purified by silica gel column chromatography in the solvents of DCM/MeOH (50:1 to 15:1). Yield: 132 mg, 0.15 mmol, 60.0 %. ^1^H NMR (400 MHz, Chloroform-*d*) δ 6.86 – 6.82 (m, 1H), 6.75 (s, 1H), 6.66 (d, *J* = 36.0 Hz, 1H), 6.37 (t, *J* = 5.4 Hz, 1H), 5.74 (dd, *J* = 6.3, 2.1 Hz, 1H), 4.46 (dd, *J* = 7.9, 4.7 Hz, 1H), 4.27 (dd, *J* = 7.9, 4.7 Hz, 1H), 3.83 (s, 3H), 3.74 (s, 3H), 3.60 – 3.49 (m, 13H), 3.49 –3.37 (m, 6H), 3.32 – 3.26 (m, 2H), 3.10 (d, *J* = 7.0 Hz, 1H), 2.99 (dd, *J* = 20.8, 1.9 Hz, 1H), 2.85 (dd, *J* = 12.8, 4.8 Hz, 1H), 2.71 (d, *J* = 12.7 Hz, 1H), 2.52 (t, *J* = 10.4 Hz, 1H), 2.20 (t, *J* = 7.4 Hz, 3H), 2.14 (s, 3H), 2.08 – 2.03 (m, 2H), 1.89 (d, *J* = 14.7 Hz, 2H), 1.81 – 1.75 (m, 1H), 1.70 – 1.62 (m, 4H), 1.46 – 1.35 (m, 4H), 1.30 (s, 3H), 1.24 (d, *J* = 3.4 Hz, 1H), 1.20 (s, 3H), 1.13 (s, 3H), 1.08 (s, 3H), 0.96 (d, *J* = 13.6 Hz, 1H), 0.67 (s, 3H).^13^C NMR (101 MHz, CDCl_3_) δ 178.22, 173.57, 164.18, 150.94, 149.69, 144.79, 144.73, 127.77, 125.54, 117.60, 106.20, 77.36, 70.47, 70.40, 70.14, 70.12, 70.04, 69.74, 61.88, 60.43, 60.35, 55.90, 55.71, 53.56, 44.43, 43.82, 40.52, 39.24, 37.51, 37.27, 36.95, 35.99, 35.03, 34.56, 34.06, 34.00, 31.70, 30.98, 30.76, 30.23, 30.06, 29.00, 28.32, 28.18, 27.97, 25.75, 23.04, 18.45, 11.95. HRMS (m/z): [M+ H] ^+^ calcd. for C_49_H_77_N_4_O_8_S, 881.5462; found, 881.5408.

**Supplementary figure source data 4.** Data for the structures, HRMS spectrum, ^1^H-NMR spectrum and ^13^C-NMR spectrum of analogue-PEG3-biotin.

Celastrol-PEG4-alkyne (*N*-(2-(2-(2-((2-propynyloxy) ethoxy) ethoxy) ethoxy) ethyl) celastrol-20-carboxamide) was generated by direct conjugation of celastrol with alkyne-PEG4-amine in DMF in the presence of BOP and TEA as previously described (Cheng et al. 2020). The product was purified by silica gel column chromatography in the solvents of DCM/MeOH (50:1 to 10:1). Yield: 180 mg, 0.27 mmol, 61.6 %. ^1^H NMR (400 MHz, Chloroform-*d*) δ 6.99 (d, *J* = 7.1 Hz, 1H), 6.48 (s, 1H), 6.37 (t, *J* = 5.4 Hz, 1H), 6.31 (d, *J* = 7.2 Hz, 1H), 4.17 (d, *J* = 2.4 Hz, 2H), 3.70 – 3.61 (m, 9H), 3.58 (d, *J* = 4.6 Hz, 2H), 3.53 (dt, *J* = 4.6, 2.9 Hz, 2H), 3.46 – 3.41 (m, 2H), 3.29 (q, *J* = 5.1 Hz, 2H), 2.46 – 2.40 (m, 2H), 2.18 (s, 3H), 2.13 – 2.07 (m, 1H), 1.98 (t, *J* = 13.4 Hz, 2H), 1.89 – 1.80 (m, 3H), 1.68 – 1.59 (m, 3H), 1.55 – 1.49 (m, 2H), 1.40 (s, 3H), 1.22 (s, 3H), 1.12 (s, 3H), 1.09 (s, 3H), 1.01 – 0.95 (m, 1H), 0.91 – 0.79 (m, 1H), 0.60 (s, 3H). ^13^C NMR (101 MHz, CDCl_3_) δ 178.35, 178.00, 170.53, 164.87, 146.07, 134.26, 127.40, 119.54, 118.09, 117.21, 79.61, 77.36, 74.77, 70.61, 70.50, 70.42, 70.13, 69.71, 69.08, 58.46, 45.14, 44.45, 43.09, 40.34, 39.42, 39.22, 38.26, 36.44, 35.02, 33.80, 33.61, 31.70, 31.14, 30.86, 30.10, 29.44, 28.74, 21.80, 18.36, 10.34. HRMS (m/z): [M+ H] ^+^ calcd. for C_40_H_58_NO_7_, 664.4213; found, 664.4166.

**Supplementary figure source data 5.** Data for the structures, HRMS spectrum, ^1^H-NMR spectrum and ^13^C-NMR spectrum of celastrol-PEG4-alkyne.

### Animal husbandry and drug treatment

Protocols for animal experiments were approved by the Committee on the Use of Live Animals in Teaching and Research (CULATR NO: 3755-15), University of Hong Kong. Obesity was induced in male C57BL/6N mice (age, 3-4 weeks; body weight, 11-13 g) by feeding on 60 kcal% HFD (Research Diets, Inc, New Brunswick, NJ, USA) for 12 weeks before the experiments. Control mice were maintained on chow diet (13.5% from fat calories) (Lab Diet, Inc, St. Louis, MO, USA). Animals were housed under 12 h of light and 12 h dark cycle with unrestricted access to food and water at Laboratory Animal Unit, University of Hong Kong. Celastrol was dissolved in saline containing 5% DMSO and 1% Tween-20 prior to administration. For drug treatment, mice were daily administrated with 5 or 7.5 mg·kg^-1^·d^-1^ celastrol or celastrol analogue via oral gavage for 21 consecutive days. Control mice and HFD mice received vehicle administration similarly by oral gavage. The body weight of mice was daily measured.

### Hematoxylin and eosin (H&E) staining

Following drug treatment, the tissues were examined by H&E staining as previously described (Cheng et al. 2015). In brief, the livers and epididymal adipose tissues were collected from diet-induced obese mice after 21-day treatment with celastrol. Tissues were fixed in 10% formalin in phosphate buffer at room temperature for at least 72 h. After sequential dehydration with 30%, 50%, 70%, 95%, 100% ethanol and xylene, the tissues were embedded in paraffin and dissected into 5 μm tissue sections. Sections were then subjected to H&E staining and examined under light microscope.

### Cell culture and drug treatment

Murine macrophage cell line RAW264.7 cells and pre-adipocyte cell line 3T3-L1 cells were obtained from the American Type Culture Collection (Manassas, VA, USA). The cells were cultured in DMEM supplemented with 10% fetal bovine serum (FBS), and 1% penicillin/streptomycin (Invitrogen, CA, USA) at 37°C in a humidified incubator containing 5% CO_2_. Binding of palmitate to BSA was carried out by a modified method as previously reported (Lu et al. 2013). Briefly, sodium palmitate was dissolved in 50% phosphate-buffered saline (PBS) and 50% ethanol to make a 100 mM stock solution. Fatty acid-free BSA was dissolved in PBS to make a solution with a concentration of 10 mg/mL. The solutions were filtered through 0.2 μm filters, mixed well to contain 50 mM palmitate, and subjected to two rounds of 10-min incubation at 55 °C towards a clear solution for use.

### Gas chromatography-mass spectrometry (GC-MS) analysis of fatty acids

Following celastrol administration, fatty acids were recovered from livers and epididymal fat pads and quantified as previously described (Quehenberger et al. 2011). Briefly, livers (50 mg) and epididymal adipose tissues (50 mg) were homogenized with stainless beads using Tissuelyzer. Organic extraction mixture containing 533 μL chloroform and 267 μL MeOH was added into 200 μL of tissue lysates (4:1 v/v) and then incubated on roller at 4 °C overnight. After centrifuging at 3000 rpm for 20 min, the lower organic (chloroform/MeOH) fraction was collected. In order to detect free fatty acid, 100 μL of organic fraction was taken to undergo methylation without saponification. The solution was firstly dried under nitrogen (N_2_) gas and then dissolved in 500 μL of 5% HCl-MeOH solution (w/v). After overnight incubation at 50 °C, the methylated lipids were extracted 3 times with 500 μL of isooctane. After dried under nitrogen gas, the extracts were dissolved in 50 μL hexane. In order to detect conjugated fatty acids, 100 μL of organic fraction was subjected to evaporation by nitrogen gas flow and saponification in 500 μL of MeOH-KOH solution at 80 °C overnight. At the end of saponification, 50 μL of concentrated HCl was added to adjust pH value to below 5.0. The lipids were extracted 3 times with 500 μL of hexane, dried by nitrogen gas flow, and methylated in 500 μL of 5% HCl-MeOH solution (w/v) at 50 °C overnight. The methylated lipids were extracted 3 times with 500 μL of isooctane, dried under nitrogen gas and dissolved in 50 μL of hexane. Following 1:10 dilution, the samples were analyzed on TRACE 1300 GC-MS system (Thermo Fisher Scientific, MA, USA) equipped with AI1310 autosampler and Thermo TG-5MS column (length, 30 m; internal diameter, 0.25 mm; film, 0.25 μm) (Thermo Fisher Scientific, MA, USA). The analysis employed several internal standards as follows: methyl palmitoleate (C16:1); methyl palmitate (C16:0); methyl γ-linolenate (C18:3); methyl linoleate (C18:2); methyl cis-9-oleic methyl ester (cisC18:1); trans 9 elaidic methyl ester (transC18:1); methyl stearate (C18:0); 5,8,11,14-eicosatetraenoic acid methyl ester (C20:4); 5,8,11,14,17-eicosapentaenoic acid methyl ester (C20:5); cis-8,11,14-eicosatrienoic acid methyl ester (cisC20:3); cis-11,14-eicosadienoic acid methyl ester (cisC20:2); cis-11-eicosenoic acid methyl ester (cisC20:1); methyl arachidate (C20:0); cis-4,7,10,13,16,19-docosahexaenoic acid (cisC22:6) and methyl docosanoate (C22:0).

### Quantitative real-time PCR determination of mRNA expression

The expression of biomarker mRNAs was determined by quantitative real-time PCR technique as described (Cheng et al. 2015). Briefly, the total RNAs were isolated from fresh liver tissues and epididymal adipose tissues using TRIzol reagent (Invitrogen, CA, USA), and converted to the corresponding cDNAs using RevertAid first-strand cDNA synthesis kit (Thermo Fisher, Waltham, MA, USA). Quantitative real time-PCR was performed with specific primers from QIAGEN (Valencia, CA, USA) and detection reagent SYBR Green mix (QIAGEN, Valencia, CA, USA). Gene-specific PCR products were subjected to melting curve analysis and quantified by the 2^-ΔΔCt^ method whereas GAPDH mRNA was determined as the internal control. The QIAGEN primers were listed as follows: adipsin (Mm_Cfd_1_SG; QT00250495), aP2 (Mm_Fabp4_1_SG; QT00091532), C/EBP-α (Mm_Cebpa_1_SG; QT00311731), CPT-1 (Mm_Cpt1a_1_SG; QT00106820), leptin (Mm_Lep_1_SG; QT00164360), PPAR-γ (Mm_Pparg_1_SG; QT00100296), resistin (Mm_Retn_1_SG; QT00093450), UCP2 (Mm_Ucp2_1_SG; QT00138943), CAT (Mm_Crat_1_SG; QT00111405), ACO (Mm_Acox1_1_SG; QT00174342), GAPDH (Mm_Gapdh_3_SG; QT01658692).

### RNA-Seq identification of differentially expressed genes

The next generation RNA-sequencing technology was employed to determine the effects of Nano-celastrol on HFD-disrupted mRNA expression as previously described (Trapnell et al. 2012). In brief, 5 mg of liver tissues were collected from three different mice in each group and homogenized in TRIzol reagent (Thermo Fisher Scientific, Waltham, MA, USA) for the preparation of the total RNAs according to the manufacturer’s instruction. The RNA quality was assessed using the Agilent 2100 Bioanalyzer (Agilent technologies, Santa Clara, CA, USA). The ribosome-depleted RNAs and polyA-enriched RNAs were prepared using TruSeq RNA sample prep kit (Illumina, San Diego, CA, USA) and subsequently converted to cDNA libraries with random hexamer primers. The library quality was assessed using the Agilent 2100 Bioanalyzer and quantified by library quantification kit (Illumina, San Diego, CA, USA). RNA sequencing was performed on Illumina HiSeq2500 platform in BGI Hong Kong (www.genomics.cn). RNA-seq data were aligned to the mouse (mm10) reference genome using STAR aligner (http://code.google.com/p/rna-star/). Differentially expressed genes was selected by DESeq2 (https://bioconductor.org/packages/release/bioc/html/DESeq2.html) using three biological replicates with a nominal significance threshold of p< 0.05 and fold change ≥ 2. Heatmaps of RNA-seq data were generated using a web-enabled heatmapper (http://heatmapper.ca/). Functional enrichment analysis of DEGs was performed online DAVID 6.8 (https://david.ncifcrf.gov/). Significantly enriched molecular function, biological process and KEGG pathway of different DEG were selected. The number of genes related to each term were shown next to the bar. Protein-protein interactions of the DEGs were analyzed by online STRING tool (https://string-db.org/). The molecular networks for DEGs in the was represented in different colors and lines using Cytoscope 3.7.2 (https://cytoscape.org/).

### Isolation of celastrol-labeled proteins

Following the treatment with celastrol-PEG4-alkyne, the cellular proteins were subjected to Click chemistry biotinylation and proteomic identification as previously described (Yang et al. 2015). In brief, RAW264.7 cells were treated with 10 μM celastrol-PEG4-alkyne for 24 h and lysed with RIPA buffer while iodoacetamide (IAA) was supplemented to block free cysteine residues in the proteins. The cellular proteins were biotinylated with biotin-azide under Click chemistry conditions (sodium ascorbate, CuSO_4_ and triazole ligand) to allow “Azide-Alkyne Cycloaddition” to occur. Biotin-labeled proteins were purified by affinity isolation on streptavidin-coated superparamagnetic beads and identified on ABI4800 MALDI TOF/TOF™ Analyzer (Applied Biosystems, Foster City, CA) through technical support for proteomics from Center of Genomic Sciences, University of Hong Kong.

### Proteomic identification of celastrol-bound peptides

Full length mouse GRP78 cDNA was cloned by RT-PCR technique using forward primer: 5’-GTCAGGATCCGAGGAGGACAAGAAGGA-3’, reverse primer: 5’-GTCACTCGAGCTACAACTCATCTTTTTCA-3’, and inserted into pET-28a bacterial expression vector after cleavage with restriction enzyme XhoI and BamHI. Competent *E. coli* BL21(DE3) cells were chemically transformed with pET28a-GRP78 plasmid DNA and induced to produce GRP78 protein by 0.2 mM IPTG at 37° C for 4 h. 6xHis tagged GRP78 was purified by HisTrap^TM^ HP column (GE Hearlthcare, Uppsala, Sweden) and HiTrap^TM^ Q HP (GE Healthcare, Uppsala, Sweden). The fractions containing GRP78 protein were combined and concentrated by centrifuging through 30-kDa Amicon ultra-15 centrifugal filter units (EMD Millipore, Billerica, MA, USA). A solution of GRP78 protein was prepared at the concentration of 6 mg·ml^-1^ in 50 mM NaHCO_3_ buffer for the binding experiments.

The binding of celastrol to GRP78 was examined at the molar ratio of approximately 1:100 for GRP78 versus celastrol. In brief, 10 µg of purified GRP78 protein was dissolved in 18 µL of 50 mM NaHCO_3_ buffer and incubated with 6 µg of celastrol in 2 µL DMSO at 4 ℃ overnight and 37 ℃ for 2 h. The protein complex was resolved by Clear-Native-PAGE (CN-PAGE) and visualized by staining with Coomassie blue. The corresponding protein band was excised as 1-mm pieces. For in-gel digestion, the gel pieces were dehydrated with 50 µL acetonitrile for 15 min and denaturated in 100 µL of 8 M urea at room temperature for 1 h. After the removal of liquid, the gel residues were washed three times with destaining buffer (25 mM ammonium bicarbonate, 50 % [v/v] acetonitrile) at 37 °C for 1 hour and subsequently dehydrated by incubating in 50 µL acetonitrile. The protein complex was digested with sequencing grade trypsin at the enzyme:substrate ratio of 1:50 (w/w) in 25 mM NH_4_HCO_3_ buffer at 37 °C for 18 h. After desalting with ZipTip C18 tips (Merck Millipore, Billerica, MA, USA), the peptides were dried by lyophilization and resuspended in 3 % acetonitrile in 0.1% formic acid.

For LC-MS/MS analysis, peptides were separated on an integrated sparytip column (100 μm i.d. × 20 cm) packed with 1.9 μm/120 Å ReproSil-Pur C18 resins (Dr. Maisch GmbH, Germany). The column was eluted with a linear gradient of solvent A (0.1 % formic acid in millipore water) and solvent B (0.1 % formic acid in acetonitrile) at the flow rate of 250 nL/min as follows: 3-7 % B at 0-2 min, 7-22 % B at 2-52 min, 22-35 % B at 52-62 min, 35-90 % B at 62-64 min, 90 % B at 64-84 min. Eluted peptides were analyzed with an Orbitrap Fusion mass spectrometer coupled with an Easy-nLC 1000 (Thermo Fisher Scientific, Waltham, MA, USA). Mass spectrometer was operated in positive polarity mode with capillary temperature of 320 °C. Full MS survey scan resolution was set to 120 000 with an automatic gain control (AGC) target value of 2 × 10^5^, maximum ion injection time (IT) of 100 ms, and for a scan range of 350−2000 m/z. The MS/MS spectra were acquired in data-dependent mode with a 3 s Top Speed method. Spectra were obtained at 15000 MS2 resolution with AGC target of 5 × 10^4^ and maximum ion injection time (IT) of 40 ms, 1.6 m/z isolation width. Raw mass spectrometry data was processed using Proteome Discover (Thermo Fisher Scientific, Waltham, MA, USA). Raw data was searched against our mouse recombinant GRP78 sequence data. Trypsin/P was specified as a cleavage enzyme allowing up to two missing cleavages. The mass error was set to 10 ppm for precursor ions and 0.02 Da for fragment ions.

### Molecular simulation of celastrol-GRP78 interactions

The crystal structure of human GRP78 (PDB ID: 3LDP) was downloaded from RCSB PDB website (https://www.rcsb.org/). The ligand molecule celastrol (PubChem ID: 122724) was obtained from the PubChem website (https://pubchem.ncbi.nlm.nih.gov/). The protein-ligand docking analysis was carried out by virtual screening software Autodock vina in PyRx 0.8 (https://pyrx.sourceforge.io/) as previously described (Cheng et al. 2020). All water molecules were removed from GRP78 protein structure prior to docking. Celstrol molecule was introduced into a 37 × 37 × 35 grid box crossing the protein structure of GRP78, automatically revealing the binding site in the protein. The docking results were analyzed and visualized by PyMOL (https://www.pymol.org/2/) and LigPlus (https://www.ebi.ac.uk/thornton-srv/software/LigPlus/).

### Western blot analysis

The protein levels of specific biomarkers were analyzed by Western blot analysis as previously described (Cheng et al. 2015, Zhao et al. 2017). In brief, 30 μg of the protein samples were resolved by 10% SDS-polyacrylamide gel electrophoresis and subsequently transferred to polyvinylidene difluoride (PVDF) membrane. Following 4 h incubation in 5% non-fat milk powder or BSA, the membranes were probed with primary antibodies against specific biomarkers overnight. The bound antibodies were detected with a goat anti-rabbit IgG-HRP conjugate. The blots were visualized by enhanced chemiluminescence (ECL) detection reagents (GE Healthcare, Uppsala, Sweden), and imaged under a Bio-Rad GelDoc imaging system (Hercules, CA, USA). The gel images were analyzed by NIH ImageJ software (http://imagej.net/ImageJ2).

### Immunofluorescence staining

The expression of specific biomarkers in livers and epididymal adipose tissues was examined by immunofluorescence staining as previously described (Luo et al. 2017, Zhao et al. 2019). In brief, livers and epididymal adipose tissue sections were firstly revitalized in retrieval buffer (pH 6.0), then permeabilized with 0.5% Triton X-100 for 30 min and blocked with 5% normal goat serum in PBS for 2 h at room temperature. Section slides were then probed with specific primary antibodies at 4 °C overnight. After 3 washes with PBS, the slides were incubated with the fluorescent secondary antibodies (e.g., Alexa Fluor 594-conjugated goat anti-rabbit IgG secondary antibody or Alexa Fluor 488-conjugated goat anti-mouse IgG secondary antibody) for 2 h at room temperature. The cell nuclei were stained with DAPI. For immunofluorescence staining of the cellular proteins, the cells were sequentially permeabilized, blocked with 5% BSA in PBS and probed with primary antibodies at 4 °C overnight. After washes with PBS, cells were incubated with the fluorescent secondary antibodies (e.g., Alexa Fluor 568-conjugated goat anti-rabbit IgG secondary antibody, Alexa Fluor 488-conjugated goat anti-mouse IgG secondary antibody and Alexa Fluor 647-conjugated donkey anti-goat IgG secondary antibody). The cell nuclei were stained with DAPI. After the removal of excessive fluorescence reagents, the liver and adipose tissues or the cells were imaged on a Zeiss LSM 780 confocal microscope (Carl-Zeiss, Jena, Germany).

### Evaluation of GRP78 chaperone activity by SPR technology

Recombinant 6xHis-tagged human GRP78 protein (Abcam, Cambridge, UK) was biotinylated with EZ-Link Sulfo-NHS-Biotin (Thermo Fisher Scientific, MA, USA) according to the manufacturer’s instructions. The biotinylated GRP78 was captured onto a streptavidin-coated Sensor Chip SA (GE Healthcare, NJ, USA) for analyzing the binding of a synthetic GRP78-targeting peptide WDLAWMFRLPVG (GL Biochem Ltd, Shanghai, China) in Biacore X100 SPR system (GE Healthcare, NJ, USA). The system was firstly equilibrated with the running buffer (10 mM HEPES, 150 mM NaCl, 3 mM EDTA, 0.005% Tween-20, pH 7.4). The binding of peptide WDLAWMFRLPVG was analyzed at the concentrations from 2.5 nM to 320 μM. In practical, the peptide in the running buffer was loaded to the system at 25 °C and a flow rate of 30 μl·min^-1^. When 10 μM celastrol was present, the peptide was analyzed at the concentrations from 1.56 μM to 100 μM. The association phase was set for 180 s, whereas the dissociation phase was set for 300 s. The sensor chip was cleaned by running regeneration buffer (10 mM glycine-HCl, pH 2.2) for 60 s before each new cycle. The affinity was determined by steady-state analysis while the estimation of Kd values was followed the 1:1 binding model with Biacore X100 Evaluation Software.

### Characterization of covalent celastrol-GRP78 conjugate

Recombinant 6xHis-tagged mouse GRP78 was expressed in *E coli* BL21 cells and purified on a HisTrap™ HP Nickel column (GE healthcare, Uppsala, Sweden). Twenty micrograms (0.3 nmol) of recombinant GRP78 was dissolved in 18 µL of 50 mM NaHCO_3_ buffer and incubated with excessive celastrol-PEG3-biotin, celastrol or celastrol analogue-PEG3-biotin at 4 ℃ overnight. The reaction mixtures were precipitated by adding cold acetone and centrifued at 13000 rpm for 15 min. The protein pellets were dissolved in 50 mM Tris buffer (pH 7.6) containing 500 mM NaCl and 1% Triton-100. An aliquote (5 µg) of protein mixtures was resolved by native SDS-PAGE and subsequently transferred to a PVDF membrane. Following overnight incubation in 5 % BSA, the membranes were probed with HRP-conjugated streptavidin (1:10000) overnight. The blots were visualized by ECL detection reagents under a Bio-Rad Gel Doc imaging system. The GRP78 was detected with anti-GRP78 antibody as index of protein loading.

### Detection of intracellular covalent celastrol-GRP78 conjugate

The formation of covalent celastrol-GRP78 conjugate was examined in RAW264.7 macrophages as previously described (Uttamapinant et al. 2012). Briefly, RAW264.7 cells were sequentially treated with 400 μM palmitate for 16 h, and 1 µM celastrol-PEG4-alkyne or celastrol for another 2 h. The cells were fixed with 4% formaldehyde in PBS (pH 7.4), permeabilized with 0.5% Triton X-100 in PBS for 30 min. Click chemistry labelling was performed by the incubation in the dark with 10 µM AFDye555-picolyl azide, 8 mM CuSO_4_, 8 mM BTTAA, and 500 mM sodium ascorbate in PBS at room temperature for 0.5 h. After washing with PBS for three times and blocking with 5% BSA, the cells were subjected to immunostaining for GRP78 and Calnexin with specific primary antibodies and fluorescent secondary antibodies. The cell nuclei were stained with DAPI. The cells were washed with PBS for 3 times and imaged under a Zeiss LSM 900 confocal microscopy (Carl-Zeiss, Jena, Germany).

### NMR determination of fat mass

The body fat contents of mice were assessed by a benchtop Bruker minispec LF90 TD-NMR analyzer from Bruker Optics Inc (Billerica, MA, USA) essentially as previously described (Luo et al. 2017). In brief, the diet-induced obese mice were treated with celastrol and celastrol analogue at the dose of 5 mg/kg/day via oral gavage for consecutive 21 days. For the determination of fat mass, the benchtop minispec instrument was firstly calibrated using Bruker standards. Mice were weighted and subsequently placed into the instrument for non-invasive determination.

### Assays of glucose tolerance and insulin sensitivity

Glucose tolerance and insulin sensitivity were assayed as described (Luo et al. 2017). For ipGTT, mice were fasted for 16 h (18:00 p.m.-10:00a.m.) and then received D-glucose at the dose of 1 g/kg. Blood samples were collected from mouse tail vein at 0, 15, 30, 60, 90, 120 min and measured for blood glucose levels using glucose test paper. For ITT, mice were starved for 6 h (10:00 a.m.-16:00 p.m.) and then received insulin at the dose of 0.75 IU/kg. Blood samples were collected from mouse tail vein at 0, 15, 30, 60, 90, 120 min and measured for blood glucose levels using glucose test paper.

### Statistical analysis

The results were presented as mean ± SD. The differences between two groups were analyzed by one-way analysis of variance (ANOVA) with Dunnett’s *post hoc* test using GraphPad Prism software (La Jolla, CA, USA). The *p*-value of less than 0.05 was considered as statistically significant.

## Acknowledgements

The authors are grateful to Ms. Lin Lin from SUSTech Core Research Facilities of Southern University of Science and Technology (Shenzhen, China) for her technical assistance in proteomic identification. The authors are grateful to Mr. Zhao Chenliang from Hong Kong Baptist University for his help in recording NMR ^1^H and ^13^C spectra of compounds. This work was supported by General Research Fund (GRF) grants (17146216, 17100317, 17119619) from the Research Grants Council of Hong Kong, the National Natural Science Foundation of China (No. 21778046) and the Seed Funding Programme for Basic Research (201611159156) from the University of Hong Kong.

## Author Contributions

Jianhui Rong designed research, Dan Luo and Ni Fan performed biochemical, cellular and animal experiments, analyzed the data and wrote the manuscript. Xiuying Zhang, Ni Fan performed peptide mapping. Fung Yin Ngo and Jia Zhao analyzed the data of RNA-seq. Wei Zhao contributed to data acquisition. Ming Huang and Yu Wang provided technical support of GC-MS analysis. Ding Li performed docking analysis. Jianhui Rong, Dan Luo and Ni Fan co-wrote the manuscript.

---

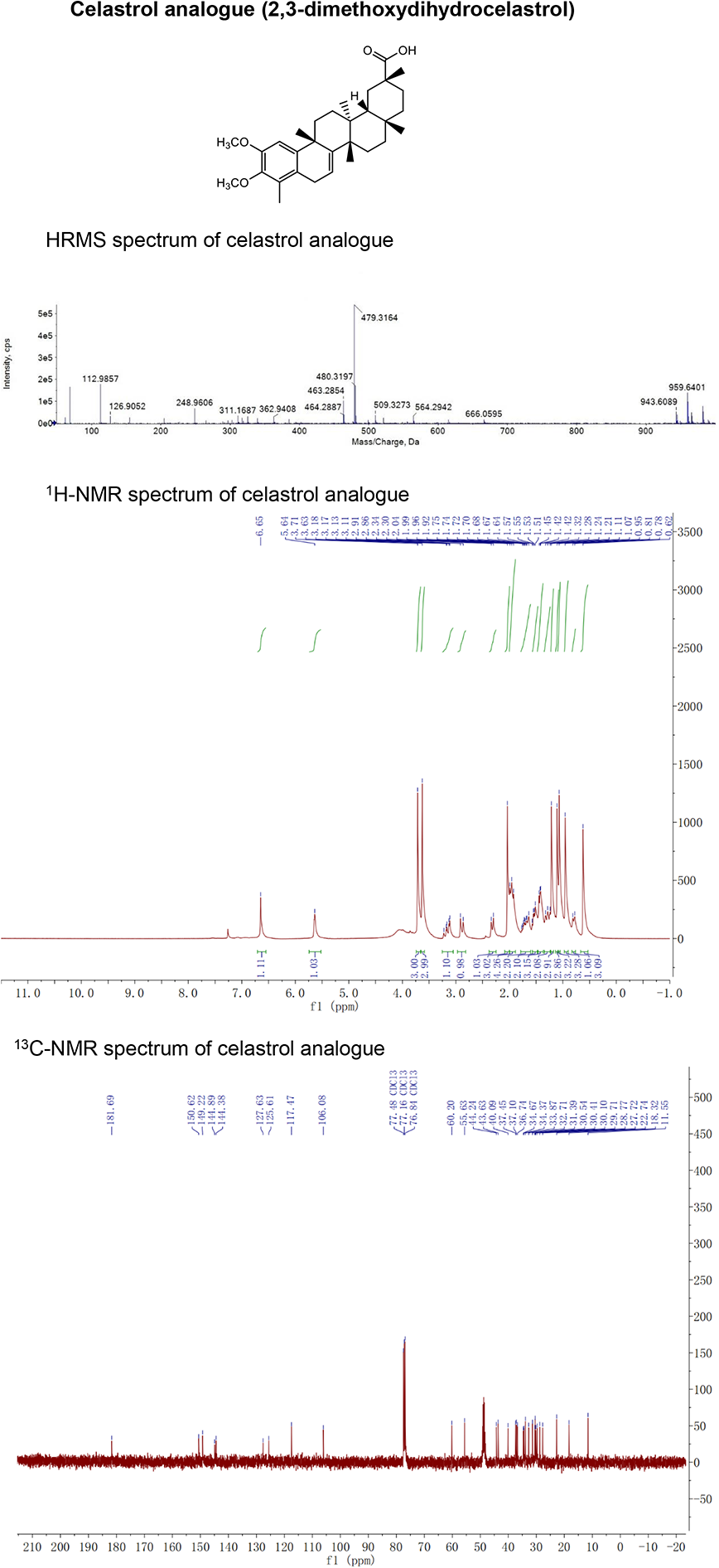

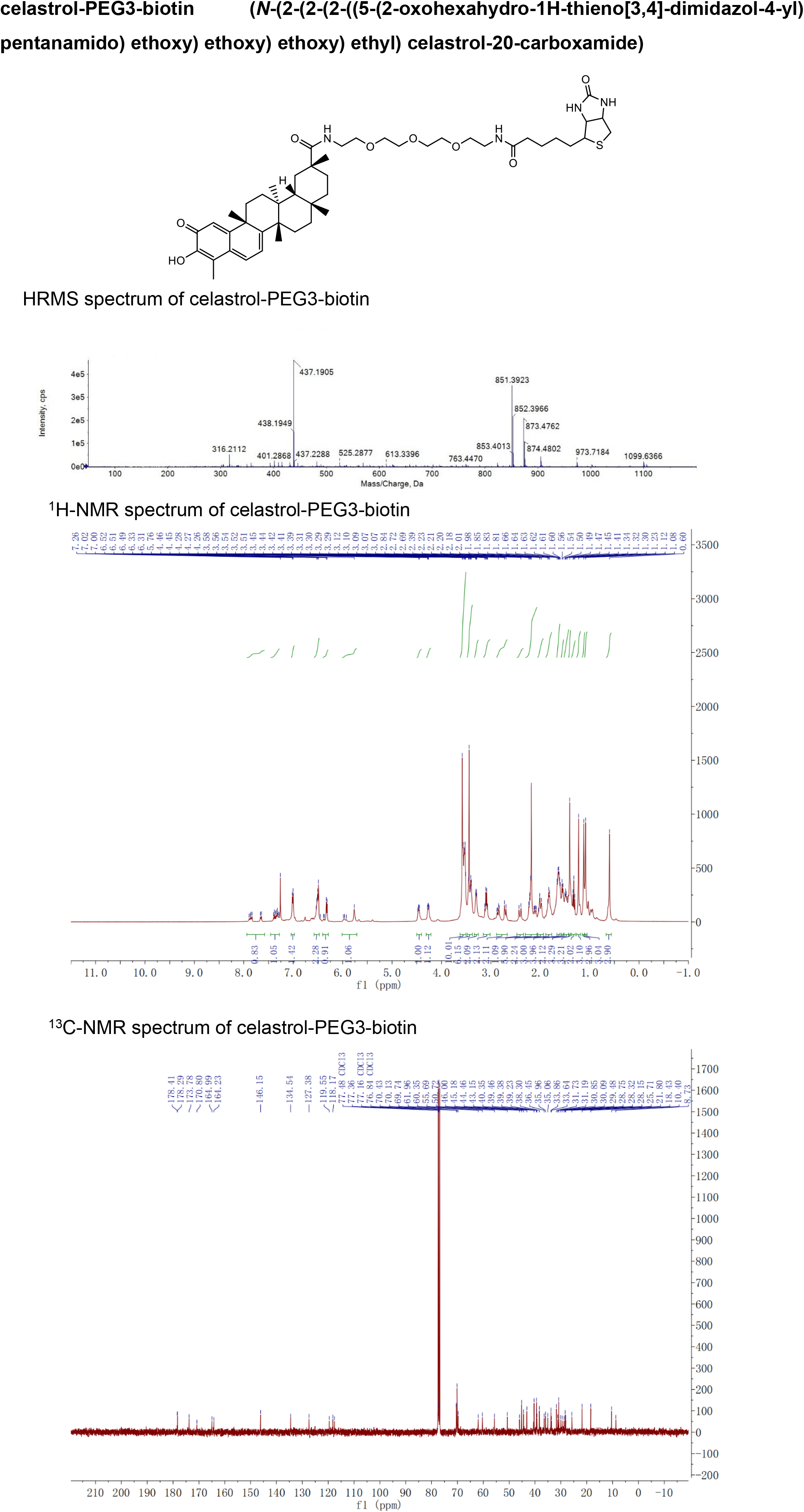

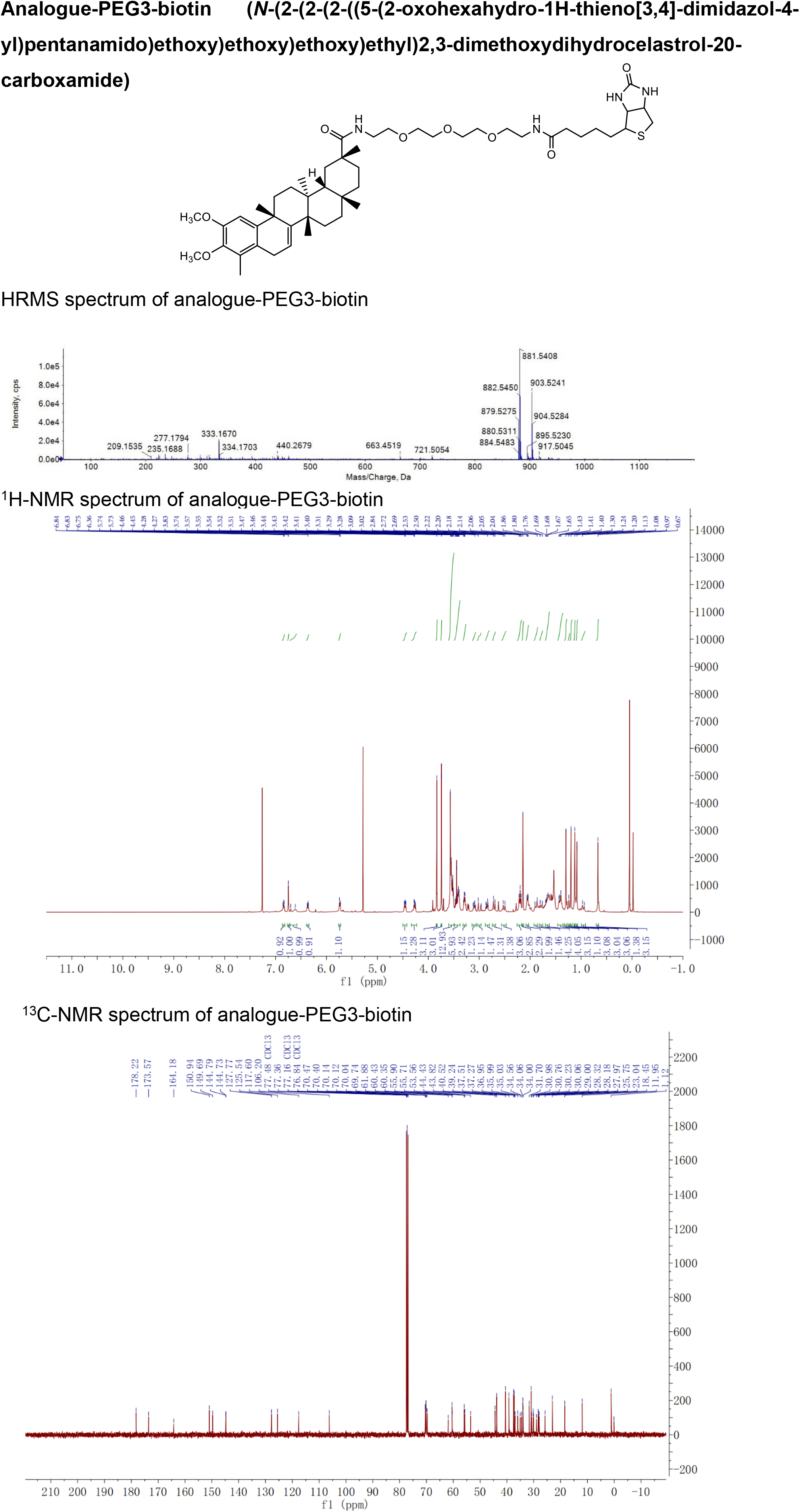

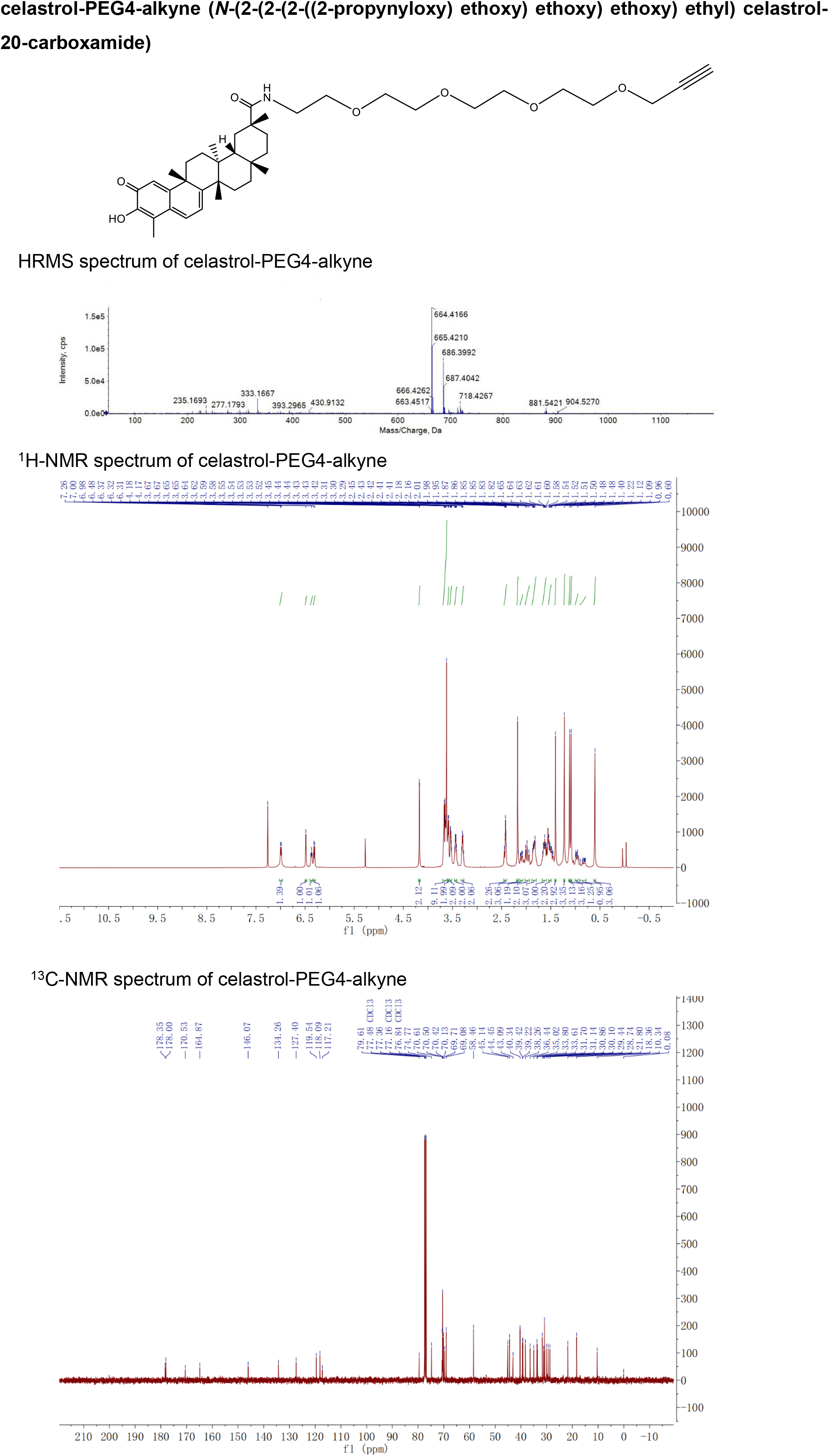

